# Interdependence patterns of multi-frequency oscillations predict visuomotor behavior

**DOI:** 10.1101/2023.04.10.536065

**Authors:** Jyotika Bahuguna, Antoine Schwey, Demian Battaglia, Nicole Malfait

**Affiliations:** Department of Psychology & Neuroscience Institute, Carnegie Mellon University, Pittsburgh, USA; Aix-Marseille Université, Inserm, Institut de Neurosciences des Systèmes (UMR 1106), 13005 Marseille, France; Aix-Marseille Université, CNRS, Institut de Neurosciences de la Timone (UMR 7289), 13005 Marseille, France; University of Strasbourg Institute for Advanced Studies (USIAS), 67084 Strasbourg, France

**Author notes:** First authorship (Jyotika Bahuguna). shared last authorship (Demian Battaglia, Nicole Malfait).

## Abstract

We show that sensorimotor behavior can be reliably predicted from single-trial EEG oscillations fluctuating in a coordinated manner across brain regions, frequency bands and movement time epochs. We define high-dimensional *oscillatory portraits* to capture the interdependence between basic *oscillatory elements*, quantifying oscillations occurring in single-trials at specific frequencies, locations and time epochs. We find that the general structure of the element-interdependence networks (effective connectivity) remains stable across task conditions, reflecting an intrinsic coordination architecture and responds to changes in task constraints by subtle but consistently distinct topological reorganizations. Trial categories are reliably and significantly better separated using oscillatory portraits, than from the information contained in individual oscillatory elements, suggesting an inter-element coordination-based encoding. Furthermore, single-trial oscillatory portrait fluctuations are predictive of fine trial-to-trial variations in movement kinematics. Remarkably, movement accuracy appears to be reflected in the capacity of the oscillatory coordination architecture to flexibly update as an effect of movement-error integration.

## Introduction

Linking neural activity to sensory, motor or cognitive processes is an ongoing goal in Neuroscience. Particular attention has been devoted to the role of brain oscillations, ubiquitous in electrophysiological recordings of both human and non-human local field potentials and EEG (Varela 2001, Buzsaki 2004). These oscillations are thought to mediate inter-regional communication (Fries 2015) and to be markers of intrinsic brain states (Fries 2015; Mantini, 2007) and task-related activity. For instance, oscillatory sensorimotor activity has been associated with motor processes decades ago (Jasper and Penfield, 1949). Since then, numerous studies have described characteristic patterns of event-related desynchronization/synchronization (ERD/ERS), calculated by averaging across single-trial absolute power time-series. Movement initiation and execution are associated with a clear decrease in beta power (beta-band ERD), and beta power typically increases (beta-band ERS) following movement offset (for review, Kilavik et al., 2013). The same power averaging procedure has been used to scrutinize finer grained relationships and to try to associate oscillatory activity in specific brain areas, frequency bands and movement time epochs with specific aspects of movement control and adaptation (Tan et al., 2014; Torrecillos et al., 2015; Arrighi et al., 2016; Savoie et al., 2018; Alayrangues et al., 2019; Jahani et al., 2020). It has been shown that theta and beta activity in medial frontal cortex are both sensitive to performance feedback and reward (Cohen et al. 2007; Marco-Pallares et al. 2007), whereas consciously perceived movement error elicits theta and beta/alpha bands responses in different cortical regions (Torrecillos et al., 2014; Alayrangues et al., 2019). Beta-band activity during movement preparation and after movement is differently modulated by movement-execution error (Torrecillos et al., 2015). Furthermore, the role of oscillations in each frequency band is contingent on the brain area where they propagate. For example, beta-band activity in medial frontal areas is involved in cognitive control of movement, while beta-band activity in lateral sensorimotor areas is modulated in relation to implicit sensorimotor adaptation (Jahani, et al., 2020).

These findings offer a glimpse of the complexity of the overall picture, but also points to limitations intrinsic to several of the evoked studies. First, searching for univocal correspondences between specific sensory, motor or cognitive processes and specific space-frequency-time oscillatory activities may well be suboptimal, and will always give only fragmentary descriptions. Second, and perhaps even more fundamentally, interpretations regarding the specific functional roles of features identified in average data may be questionable because these features may not even exist in the individual trials. Typically, the slow fluctuations (ERD/ERS) visible in trial-averaged power profiles cannot be detected in individual trials (e.g. Tallon-Baudry & Bertrand, 1996; Naik et al., 2021); they do emerge as new and abstract features from smoothing out inter-trial fluctuations, treated as uninformative noise. But, concretely, behavior is performed in each individual trial, and also varies from trial to trial. Hence, variability in single trial oscillatory activity is not only mere noise, and its analysis may reveal neural mechanisms that are not apparent in cross-trial averages. Driven by this idea, previous studies could successfully correlate inter-trial behavioral and electrophysiological variations using linear models on single trial data (Cohen & Cavanagh, 2011; Torrecillos et al., 2018; Lofredi et al., 2019). However, these studies still suffer from the limitation we point out above, as they focus on oscillatory activity observed in a given brain region, frequency band and trial epoch, and thus offer fragmentary views. It is unlikely that sensory, motor or cognitive operations are implemented and revealed by individual, wildly fluctuating oscillatory activity, localized in space, time and frequency.

Oscillations at different locations, times, and frequencies are likely to be comodulated by modes of system’s level collective dynamics, rather than constituting collections of completely independent processes (Atasoy et al., 2016). It has been proposed, for instance, that oscillations can be used to selectively gate information transfer between neuronal populations even when they are irregular and stochastic, as long as their irregular fluctuations coordinate across time, space and frequency (Palmigiano et al., 2017). Such coordination arises in a self-organized manner, as the system is constrained by global structural and dynamical determinants to sample lower-dimensional manifolds within the high-dimensional space of configurations it could theoretically access if its parts were separately controlled (Bressler & Kelso, 2001; Pillai & Jirsa, 2017). Structured behavior along a task would thus just be associated to transient adaptations of system’s ongoing trajectories on these manifolds, rather than to abrupt reconfigurations (Shine et al., 2019; Naik et al., 2021). These dynamical system views are akin to early proposals that evoked activity and networks are very similar to spontaneous fluctuation patterns (Kenet et al., 2003; Cole et al., 2014, Naik et al., 2021), possibly reflecting capability for probabilistic computations (Orban et al., 2016).

Here, we hypothesized that the fluctuations of single-trial oscillatory *elements* (power at a given space-frequency-time point) are tightly coordinated and that the structure of their interdependence network reflects an intrinsic coordination architecture that responds to changes in task constraints by subtle but nevertheless consistent topological reorganizations. To assess this idea, we analyzed EEG signals recorded during a motor adaptation task representative of the richness of possible oscillatory behaviors and functional mechanisms of cognitive and automatic movement control and monitoring (see Jahani et al., 2020). To capture the collective dynamics of the elements in a global state space, we defined high-dimensional single trial oscillatory *portraits* containing all individual oscillatory elements of a given trial. In agreement with our hypothesis, we found that each single-trial oscillatory element could reliably be predicted based on the knowledge of the other ones. Hence, the fitted models described a form of “effective connectivity” (EC; Friston, 1994; 2011) between the oscillatory elements, quantifying mutual directed influences, beyond simple correlations. These networks of interdependencies between oscillatory elements thus provide the ultimate description of the complex patterns of coordinated dynamics underlying the implementation of behavior.

We found that the EC between oscillatory elements had a general structure that remained stable across the different task conditions, likely to be intrinsic (Friston 1994) and reflect dynamical constraints imposed by structural connectivity (Mostame et al., 2021, Honey et al., 2007). However, the EC network exhibited fine adaptations in their detailed topology in response to changes in the task constraints, as a potential mechanism underlying rich and flexible task-condition-specific behavioral adjustments. Trials performed in the different task conditions could be reliably distinguished based on these specific EC network reorganizations, detectable through the observation of fluctuating single trials portraits. In addition, we found that these portrait fluctuations were also coupled with fine trial-to-trial variations in movement kinematics. For instance, movement error directly modulates the degree of interdependence between oscillatory elements. In other words, the EC network is fine-tuned as an effect of integrating movement error information, with a deficit in this fine-tuning being associated with unsuccessful behavior.

Altogether, our findings suggest that visuomotor cognition and behavior are supported by collective brain states regulating the integrated dynamics of distributed sub-systems, rather than by a multitude of segregated oscillatory processes.

## Results

### Capturing inter-trial variability of single-trial elements via oscillatory portraits

EEG was recorded in volunteers performing a motor adaptation task in which they had to “shoot”, without stopping, a visual target, making ballistic reaching movements (Figure 1A). When successful, the target exploded making the sound of a bottle of champagne being uncorked. The target was small enough to make it not trivial to touch it, and participants indeed missed it about half of the time (this, under baseline usual conditions – *NR* trials). Moreover, making the task even more challenging, the visual feedback of the hand (a cursor) was rotated relative to its actual position for short series of 4 trials (Figure 1A). The visual rotation was unexpectedly (re)introduced, so that on the first trial (*Ct* trials) of each “rotation-trial-series” participants saw the cursor going in the wrong direction, and they systematically missed the target. Nevertheless, as soon as they knew that the visual rotation was turned on, they were able to counter it by applying a cognitive strategy. As a matter of fact, before the experimental session, we explained them in detail the nature of the visual perturbation, and how they could exactly compensate for it by aiming at a neighboring target (*RS* trials). Still during these trials, the discrepancy between the visual and proprioceptive information activated automatic updating of the sensorimotor map (implicit sensorimotor adaptation), which was testified by the slight movement deviations (“after-effects” in the direction opposite to the visual rotation) visible upon the removal of the perturbation (*AR* trials).

**Figure 1:**
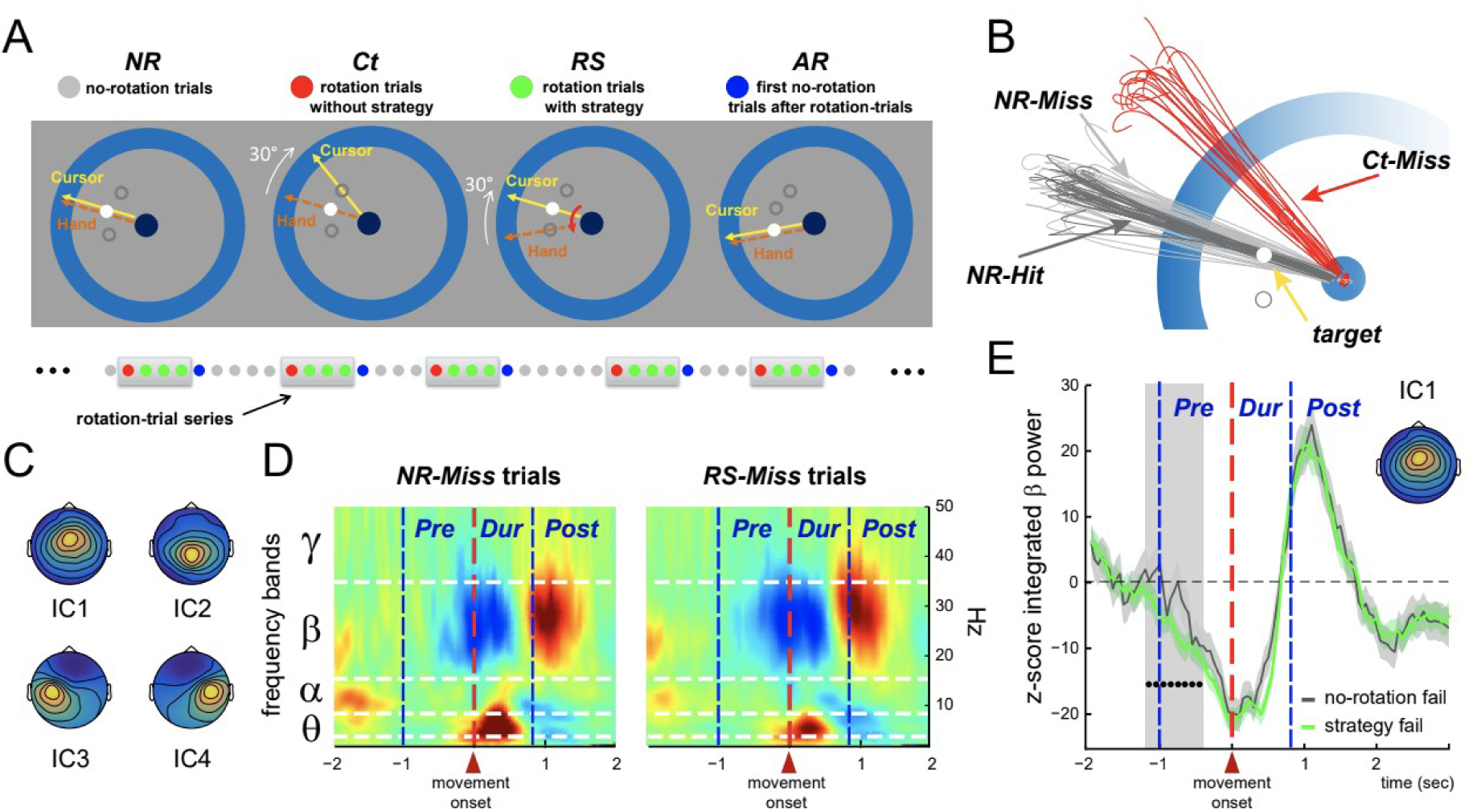
Analyzed data: independent component (IC) time-courses. **A)** EEG was recorded in healthy participants instructed to “shoot,” without stopping, one of three possible visual targets. In Ct (red) and RS (green) trials, the cursor representing the index fingertip was displayed rotated by 30° relative to its real position. In the Ct trials the rotation was unexpectedly introduced and thus produced large kinematic errors. In contrast, in the RS trials, participants applied a strategy to counter it. These rotation trials were separated by trials without visual rotation: one AR (blue) trial and a variable number of NR (gray) trials. **B)** Trials were further divided into Hit and Miss, as illustrated by individual hand paths by one participant. **C)** Group average topographies of the four ICs of interest, capturing respectively activity in the frontal medial cortex (IC1), the parietal medial cortex (IC2), and the anterior and posterior blanks of the central sulcus, left (IC3) and right (IC4). **D)** Group average time-frequency representations of the activity of IC1 in the NR-Miss (left) and the RS-Miss (right) trials). IC time-courses are aligned to movement onset. The different trial epochs (Pre, Dur, Post) are indicated by the vertical dashed lines. The different frequency bands (θ, α, β, γ) are indicated by the horizontal dashed lines. **E)** Group average beta-power profiles for IC1 aligned to movement onset, for the NR-Miss (gray) and the RS-Miss (green) trials respectively. The black dots indicate the period during which beta-power was significantly lower in RS-Miss trials relative to NR-Miss trials.

In our analyses, we distinguished between the trials in which participants successfully shot the target (*Hit*) from those in which they failed to do so (*Miss*). We considered the following 6 categories of trials: *Ct-Miss*, *RS-MIss*, *RS-Hit*, *AR-Miss*, *NR-Miss* and *NR-Hit*.

The signal recorded by an EEG electrode applied on the scalp is a mixture of multiple signals arising from different sources, neuronal or not. Here, we used temporal independent component analysis (ICA) as a blind source separation technique and identified in each participant four independent components (IC) capturing respectively oscillatory activity of the following four cortical areas: the frontal medial cortex (IC1), the parietal medial cortex (IC2), and the anterior and posterior blanks of the central sulcus, left (IC3) and right (IC4). Each IC is characterized by a time-invariant topography (spatial filter) and an activation time-course. Figure 1C presents the topographies of the four ICs (obtained by averaging the topographies of the IC identified for each participant). The activation time-courses of the ICs can be subjected to the same analyses as usual EEG signals. Figure 1D presents group-average (across trials and participants) spectrograms of the time course of IC1 (frontal medial region) for two different categories of trials, *RS-Miss* trials (right) in which participants applied the cognitive strategy to counter the visual rotation and *NR-Miss* trials (left) in which they did not have to, as the visual display was not altered (see Figure 1A). Overlaid on the spectrograms, the horizontal and vertical dashed lines indicate respectively the frequency bands (θ, α, β, γ) and the movement periods (*Pre*, *During*, *Post*) of interest. We originally analyzed the present EEG data set using conventional trial averaging procedures (Jahani et al., 2020). We found that beta-band activity of the frontal medial cortex was selectively attenuated during movement planning when participants applied the re-aiming strategy. Figure 1E shows the group-average beta power profile of IC1 (frontal medial cortex) for the *RS-Miss* and *NR-Miss* trials. One can see that beta power is significantly decreased during the *Pre* trial-epoch for the *RS-Miss* trials, in which participants applied the cognitive strategy, relative to the *NR-Miss* trials, in which they did not have to, as there was no visual rotation. Similar analyses has been conducted for power at other possible combinations of brain location, frequency and time period (Tan et al., 2014; Torrecillos et al., 2015; Alayrangues et al., 2019).

However, interpretations regarding the specific functional roles of features identified in averaged data may be questionable because these features may not even exist in the individual trials. Here, the slow fluctuations (ERD/ERS) clearly visible in trial-averaged time-frequency maps (Figure 1D) do not exist at the single trial level (Figure 2A); they are in fact subtended by short-lived oscillatory bursts (Murthy et al., 1992; Tallon-Baudry et al., 1996; Feingold et al., 2015) and do emerge as new features from smoothing out inter-trial fluctuations. We can provide a coarse-grained description of each of these single-trial spectrograms by averaging power over specified trial epochs (*Pre*, *During* and *Post* movement) and different frequency bands (θ, α, β, γ). Each of these averages provides the value of what we call an *oscillatory element* of the trial, and quantifies power observed at one of the brain regions (IC1, IC2, IC3 or IC4), in one of the frequency bands (θ, α, β, γ) and during one of the trial epochs (*Pre*, *During* and *Post*). This results in 48 oscillatory elements –one for each possible combination of space, frequency and time– for each trial (see Figure 2B).

**Figure 2:**
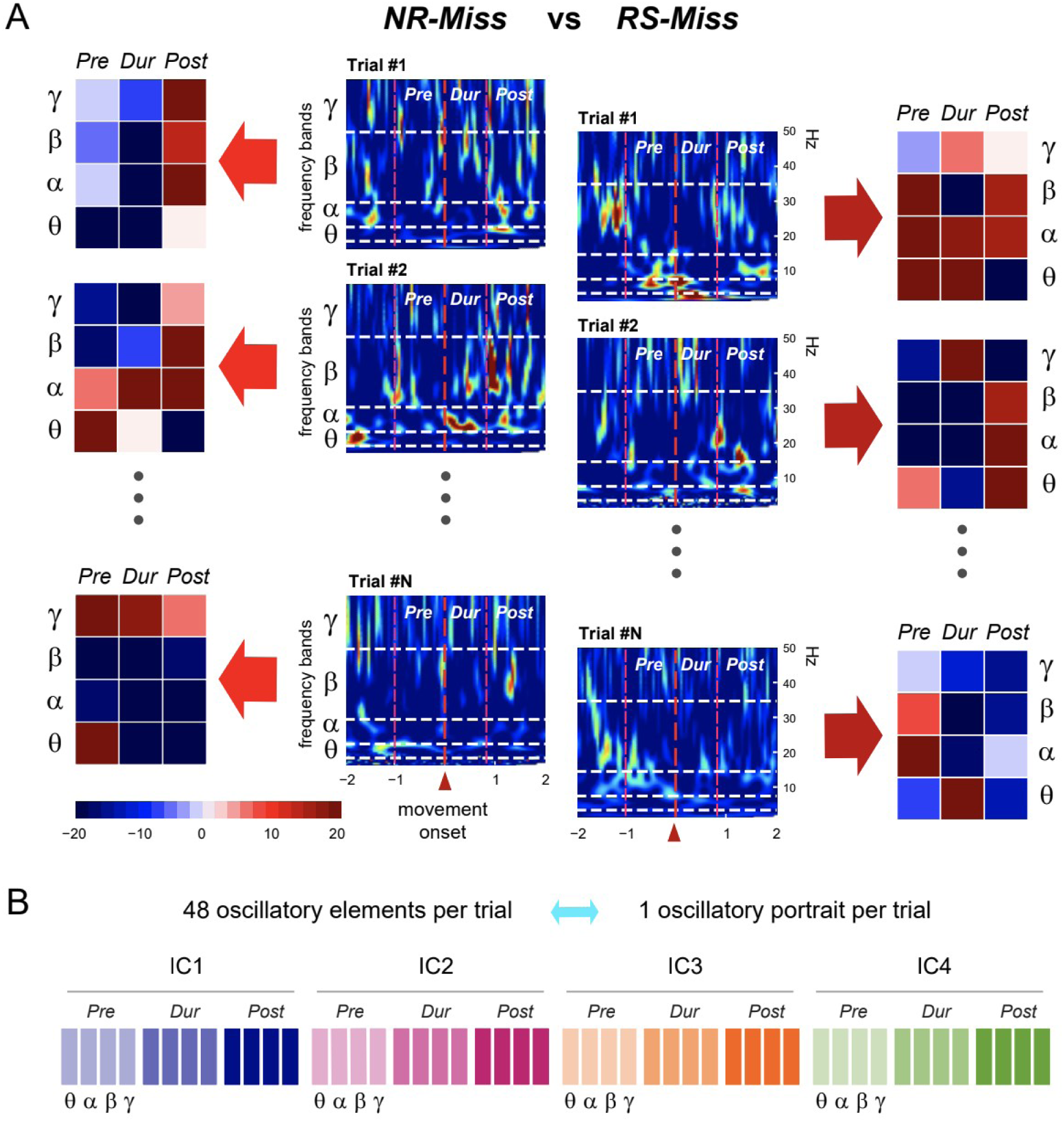
Oscillatory elements and portraits. **A)** Single-trial activity of IC1 (frontal medial cortex) in the NR-Miss and the RS-Miss trial categories. From each single-trial time-frequency map (center columns), oscillatory elements are obtained by averaging power within each trial epochs and frequency bands (left and right columns). **B)** Each oscillatory element quantifies single-trial power at a given brain location (IC1, IC2, IC3, IC4), trial epoch (Pre, During and Post movement) and frequency band (θ, α, β, γ). Single-trial portraits are composed of all 48 (4 ICs x 3 trial epochs x 4 frequency bands) single-trial elements.

To capture the potential interdependence between the single-trial oscillatory elements, we combined them into 48-dimensional vectors providing synthetic and multivariate characterization of each trial, which we named single-trial *oscillatory portraits* (see Figure 2B). The values of the oscillatory elements associated with the illustrative trials in Figure 2A are represented within colored matrices (portrait chunks) next to the corresponding single-trial spectrograms. Even if the coarse-graining in time and frequency smooths out some of the variability, the wide intertrial differences are still visible from these element matrices. As a result of this variability, the ranges of values observed for a given oscillatory element in the different categories of trials largely overlap. We can precisely define this degree of overlap by evaluating the probability that single-trial values of a considered element also fall within a common reference range of fluctuation (*Pre*-period for success trials, see *Materials and Methods*). As shown by Figure S1, these overlaps were always above ∼80%. How do these large overlaps in the distribution lead to statistically significant differences between trial categories at the level of average comparisons?

To compute averages and compare them it is necessary to have access to many observations of fluctuating oscillatory elements, to sufficiently sample their distribution. However, the question we want to ask here is whether an observer having access to just *one* observation of an oscillatory element could successfully infer (or not) the category of the trial within which the considered element has been observed. Such an observer (that could represent, for instance, a machine learning classifier attempting trial category discrimination, or even another reader neuronal population, analyzing oscillatory activity during actual behavior), to perform the inference, should compute the likelihood that the observed element comes from one or the other distribution, associated to different trial categories. However, these likelihoods would be very similar for most trial categories, as the probability overlap between distributions is so large. Because of these overlaps, the inverse problem of inferring which distribution the observed element was sampled from becomes thus extremely difficult, if not impossible, to solve.

We could however formulate the hypothesis that the fluctuations of individual oscillatory elements are not independent (as expected in a scenario in which collective system’s dynamics is jointly comodulating many elements). In this case, fluctuations would not be suppressed but would occur over restricted manifolds of element interdependence and these manifolds may be slightly different for different categories of trials and behaviors. An observer could thus discriminate which type of behavior has occurred by decoding over which manifold the observed oscillatory portrait (fluctuating in many dimensions) is located. Individual elements could still fluctuate over overlapping ranges, as the different manifolds specific for different trial categories would have overlapping projections over the unique dimension of variation of each element. However, in the high-dimensional space of the entire oscillatory portraits, the different manifolds could still be distinguishable.

To investigate the plausibility of this hypothesis we proceeded in two steps. First, we verified that, indeed, elements were inter-dependent between them and that, therefore, the fluctuations of oscillatory portraits did occur on lower-dimensional subspaces of their overall 48-dimensional space (Figure 3). Second, we showed that the subspaces of co-fluctuation were not identical for different trial categories (Figure 4), so that a classifier can learn to to distinguish them, when inspecting whole oscillatory portraits rather than individual oscillatory elements one at a time (Figure 5).

**Figure 3:**
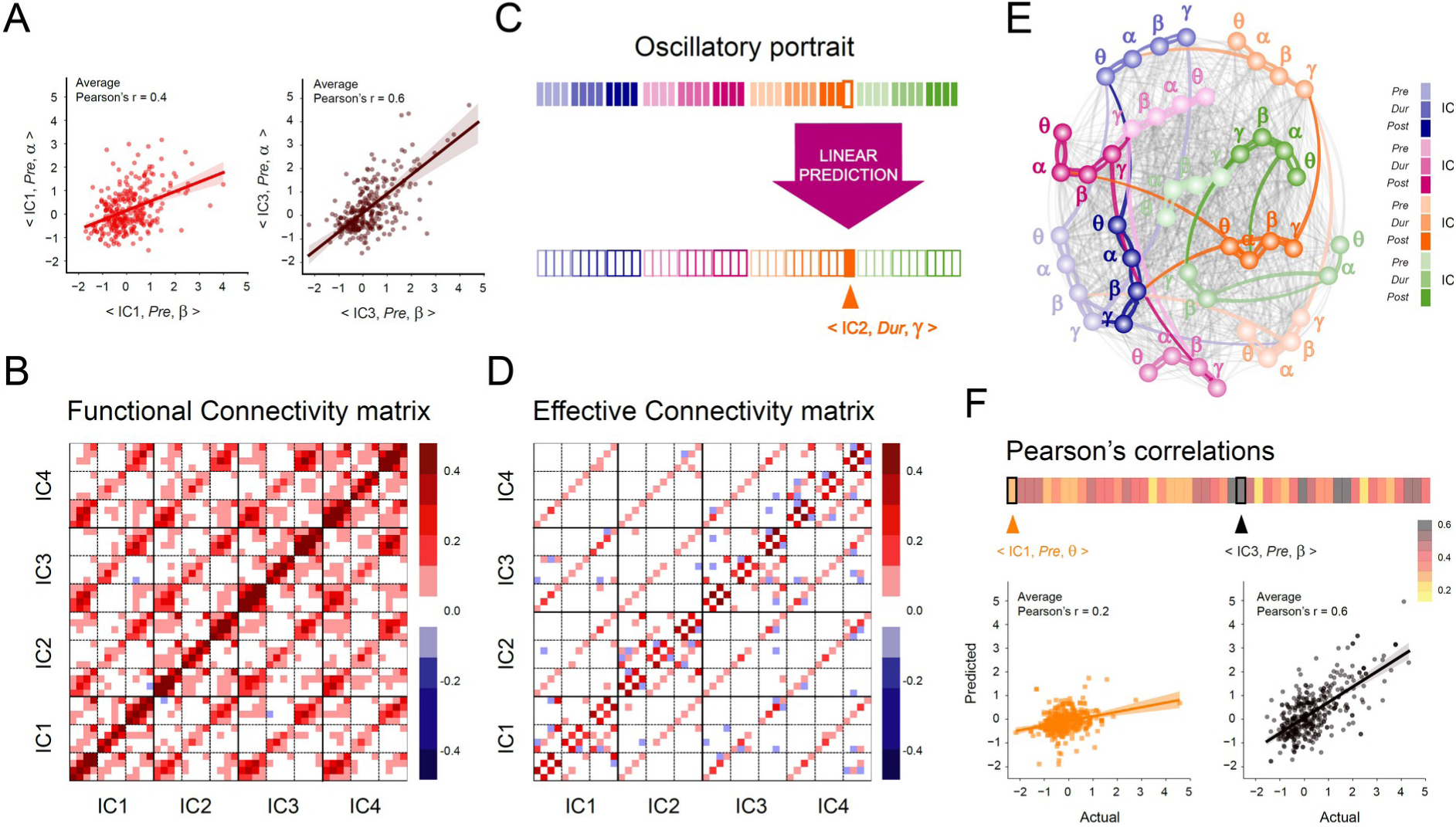
Effective connectivity between oscillatory elements. **A)** Individual spatio-temporo-spectral oscillatory elements may show strong covariation. Illustrative scatter plots for two pairs of oscillatory elements (< IC1, Pre, α > vs < IC1, Pre, β > and < IC3, Pre, α > vs < IC3, Pre, β >) showing significant covariation. **B)** The correlation matrix (functional connectivity) reveals mostly positive values. **C**) To quantify effective connectivity, multivariate linear regression was performed to predict the values of each single-trial element based on those of the 47 other entries of the same single-trial oscillatory portrait. **D)** The effective connectivity (EC) matrix is sparser than the correlation matrix, but still shows dependence across all spatial locations. **E)** Graph representation of the effective connectivity matrix in D. The nodes represent individual oscillatory elements and the edges figure the effective connectivity links between them. The spatial locations (ICs) are indicated by colors and trial epochs by color shades. The nodes are labeled according to the frequency band. Strong connections are shown as thick lines colored according to the source nodes. Thin gray lines show weak connections. **F)** The single-trial oscillatory elements could be predicted from others with varying degrees of accuracy. Scatterplots of actual against predicted values are presented for two elements, < IC1, Pre, θ > and < IC3, Pre, β >, for which average Pearson’s correlation ∼ 0.2 and ∼ 0.6, respectively.

**Figure 4:**
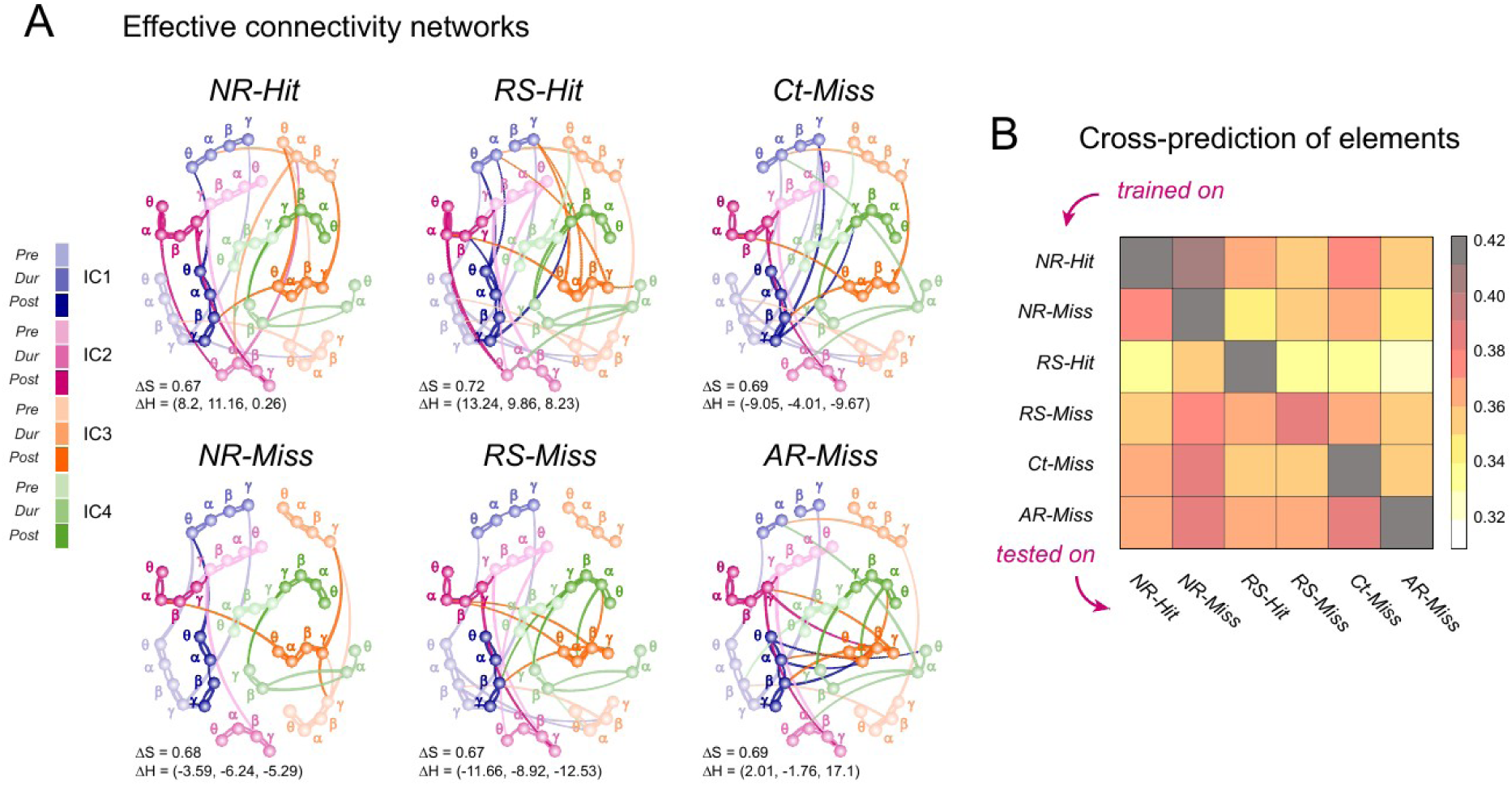
Similar but distinct effective connectivity (EC) networks for the different trial categories. **A)** Graph representations of the EC for each of the 6 trial categories separately. The nodes represent individual oscillatory elements and the edges figure the effective connectivity links between them. The spatial locations (ICs) are indicated by colors and the trial epochs by color shades. The nodes are labeled according to the frequency. Strong connections are shown as thick lines colored according to their source nodes. Weak connections are not shown. **B)** The similarities between the EC networks specific to the different trial categories were quantified by the achieved cross-prediction performances (see Material and Methods); that is, how well a classifier trained on trials of one given category predicts the values of the single-trial oscillatory elements of trials of a different category. Prediction performance was measured by the average Pearson’s correlations between the actual and the predicted values of the oscillatory elements. The lowest Pearson’s correlations were around 0.3

**Figure 5:**
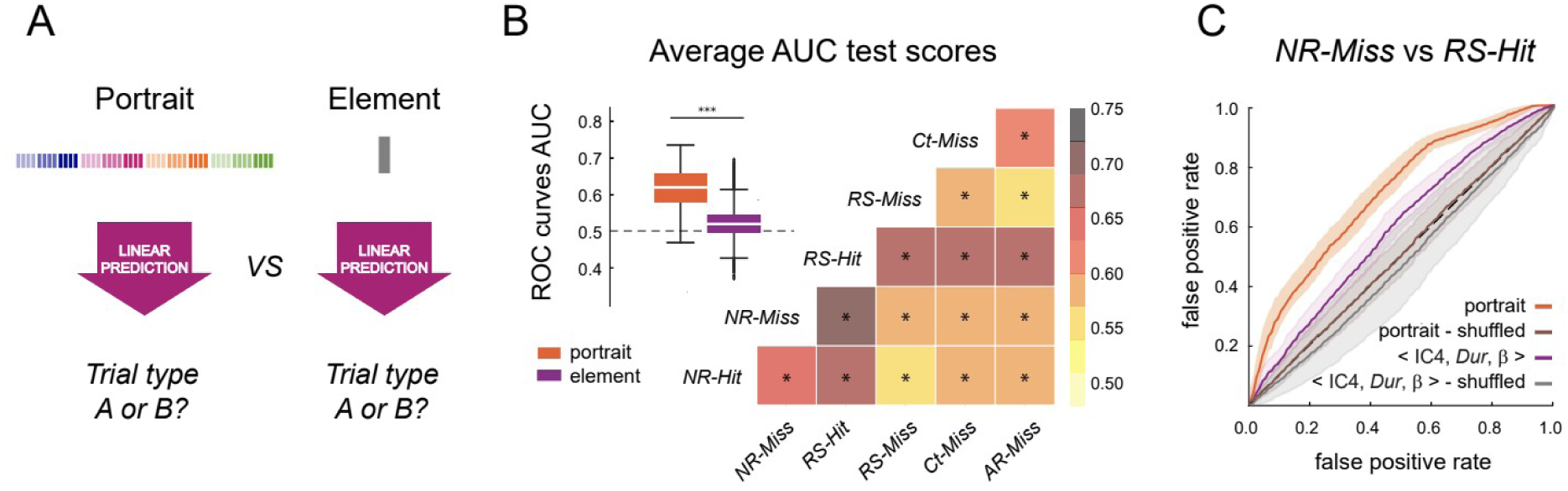
Trial categories are reliably separated by classifiers trained on full portraits. **A)** We compared the accuracy of category pairwise separations achieved by portrait-based and element-based classifiers. **B)** Separation performance was measured by the area under the curve (AUC) for ROC curves. On average (boxplots) portrait-based classifiers performed with > 60% accuracy, whereas element-based classifiers yielded marginally above chance level (0.52 vs 0.5, two-sided sample t-test: 69.9, p=0.0) performances. Matrix summarizing the average AUC for each trial-category pairwise-separation achieved by portrait-based classifiers; in all cases, AUC was significantly above chance level (0.5). **C)** For illustration, ROC curves for the separation of the NR-Miss and RS-Hit trials by classifiers trained on full portraits versus single elements and shuffled data labels. The classifier trained on full portrait outperforms (0.71± 0.03) the classifier trained on the most predictive single element < IC4, Dur, β > (0.59 ± 0.02).

### Oscillatory elements within portraits are functionally and effectively connected

As a first step, we studied the correlations between the fluctuations of the different oscillatory elements. Each single-trial oscillatory portrait was considered as a single point within a high-dimensional (48-D) space and we sought for structured covariances within the cloud of points formed by all of them. Figure 3A shows two representative scatter plots of one oscillatory element vs another one: < IC3, *Pre*, α> versus < IC3, *Pre*, β > on the left, and < IC1, *Pre*, α > versus < IC1, *Pre*, β > on the right. These two scatter plots display markedly positive linear correlations between alpha and beta powers in the left sensorimotor (IC3) and medial frontal (IC1) cortices during the pre-movement period (significant Pearson correlation values: ∼0.4, p = 0.0003 for IC1 and ∼0.6 for IC3, p = 0.0001). Figure 3B displays the complete correlation matrix between the 48 oscillatory elements. Besides the two examples of Figure 3A, many other pairs of elements were positively correlated (correlation matrix averaged over various bootstraps in Figure 3A - see Materials and Methods). Correlations between different frequency bands were particularly strong within brain regions (ICs) and trial epochs. Nevertheless, significant positive correlations (and a few rare negative correlations) were also found between elements of different ICs and trial epochs. The observed correlation matrix was thus rather dense, confirming that the different oscillatory elements are linked by a *functional connectivity (FC)*; that is, they fluctuate in a coordinated manner.

In a second step going beyond correlational observations, we attempted to predict the fluctuations of one oscillatory element based on the fluctuations of the others within the same portrait. Prediction requires the choice of a model; for the sake of simplicity, we chose simple Linear Mixed Models. Such a choice corresponds to modelling the manifolds of element covariation as hyperplanes within the 48-dimensional space of portraits. Such hyperplanes may be differently oriented for different trial categories. However, we initially studied the general element interdependence structure combining observation from trials of any type together. We thus performed multivariate regression to predict the values of each single-trial element based on those of the 47 other entries of the same single-trial oscillatory portrait (Figure 3C; see *Materials and Methods*). The resulting matrix of model coefficients is shown in Figure 3D. Such matrix of *effective connectivity* (EC) –as now directed and predictive (Friston, 1994)– revealed patterns of interdependence that are reminiscent of motifs within the correlational FC matrix of Figure 3B. The EC matrix was however much sparser.

We can adopt an alternative representation for these EC connections, in which each of the 48 oscillatory elements corresponds to a network node. In Figure 3E, the nodes are colored according to the corresponding IC, with darker or lighter hues according to the trial epoch and are labeled according to the frequency band. Strong directed links of EC directed influence are represented by thick lines colored according to the source node, while weaker ones are shown in light gray. We evaluated how well the single-trial values of an element could be predicted based on the values of the other elements of the same single-trial portrait. We used a cross-validation approach in which prediction performance is assessed on data that were not used to fit the model (see *Materials and Methods*). Figure 3F summarizes the achieved performance for each of the 48 oscillatory elements, together with the scatter plots of the actual against the predicted values for two representative oscillatory elements. The cross-validated average values ranged from ∼0.2 (for element < IC4, Dur, θ >) up to ∼0.7 (for element < IC3, Pre, β >). These results support our intuition that oscillatory portraits are entities characterized by a strong internal interdependence from which individual element fluctuations can be reliably predicted.

Although the web of EC links was dense, the strongest connections formed characteristic clusters that tended particularly to link together oscillatory elements of the same region (IC), trial epoch and frequency-band. Such visual impression was confirmed by quantitative analyses of a network connectivity feature known as *homophily* (McPherson et al. 2001). A network is considered homophilic if nodes of a certain type tend to connect with nodes of a similar type with a probability greater than chance. In our network, every oscillatory element node had three types of labels: a space label (from which IC it was recorded); a time label (from which task epoch); and a frequency label (at which frequency band). Testing for homophily corresponds thus to verifying whether the strength of connections between nodes with a same (or different) label values is enhanced (or reduced) with respect to a null model in which labels have been shuffled across nodes but connectivity wiring was maintained (see *Materials and Methods)*. We thus performed in Figure S2 separate quantifications for the three types of space, time and frequency homophily. In Figure S2A, addressing space homophily, the two lines show the average weight with which an element measured at an IC is connected with elements at another IC (independent of the trial epoch and frequency band of the considered source and target elements), for the actual EC network and surrogate space label-shuffled networks. Figure S2A reveals a marked spatial homophily (73% of the total weights are contained in spatially homophilic nodes), as there are peaks for connection strength between nodes within the same IC and, for connections between nodes at different ICs, the connectivity strength remains well below chance-level from the surrogate model. Similarly, time (85%) and frequency (62%) homophily was observed (Figure S2B and S2C respectively), even if frequency homophily was less precise, with the appearance of weaker peaks of enhanced cross-frequency connectivity, possibly reflecting cross-frequency couplings (Canolty et al., 2010). Therefore, the observed network of EC interdependency is neither homogeneous, with oscillatory elements equally connected independently from their labels, nor extremely specific with connections strictly confined with different regions, time epochs and frequency bands completely independent from each other. It is homophilic, a network configuration between complete order and complete disorder, indicative of the complexity of the oscillatory inter-dependence architecture (see *Discussion*).

### Effective connectivity between oscillatory elements is similar yet unique for different trial categories

We have demonstrated that individual oscillatory elements fluctuate in a coordinated manner and identified the structure of the EC network estimated from the trials of all the different categories pooled together. The question that arises then is whether and how this pattern of interdependence (and the corresponding co-fluctuation hyperplane in the space of oscillatory portraits) is altered by changes in the task conditions. To investigate this question, we repeated the EC fitting and model-performance quantification steps described above, separately for each category of trials. Figure 4A presents the EC networks for the six trial categories, *NR-Hit*, *NR-Miss*, *RS-Hit*, *RS-MIss*, *Ct-Miss* and *AR-Miss*. The different networks were highly similar between them and with the general EC fitted over all trial categories pooled together, in that they all exhibited strong homophily, as shown by prominent clusters of strong connections between nodes of the same region (IC), frequency band and trial epoch (Figure 3E). We quantified the similarity between the networks by measuring the Pearson Correlation CC of their adjacency matrix with the one of the pooled EC. All these correlations were significant and of the order of CC=∼0.7 (*AR-Miss*: 0.69, Ct-Miss: 0.69, NR-Miss: 0.68, RS-Miss: 0.67, NR-Hit: 0.67, RS-Hit: 0.72), denoting large overlap. Finally, we also compared relative variation in the three kinds of homophily ΔH% (see *Materials and Methods*) with trial categories as compared to the pooled EC. Interestingly, we found a decrease in spatial, spectral and temporal homophily for three *Miss* trial categories: *Ct-Miss*: (−9.05%, −4.01%, −9.67%); *NR-Miss*: (−3.59%, −6.24%, −5.29%) and *RS-Miss*: (−11.66%, −8.92%, −12.53%). However, in contrast, we find an increase in spatial, spectral and temporal homophily for the two *Hit* categories: *NR-Hit*: (8.2%, 11.16%, 0.26%) and *RS-Hit*: (13.24%, 9.86%, 8.23%). An increase in the homophily for successful trial categories may indicate an overall increase in structure, whereas decrease in the homophily may indicate an overall increase in randomness of the EC matrix. The *AR-Miss* however showed a mixed trend with an increase in spatial (+2.01%) and temporal (+17.1%) homophily but a decrease in spectral (−1.76%) homophily.

Are these subtle differences sufficiently distinctive to separate the specific manifold of element co-fluctuation for different trial categories? A model trained on trials of category *A* can be tested on trials of category *B* to assess how well its prediction performance generalizes from *A* to *B* (see *Materials and Methods*). The more similar the underlying statistical interdependence structure for trial categories *A* and *B*, the better the prediction generalizes. Such cross-training prediction approach is not dissimilar in spirit from *representational similarity analysis* (Kriegeskorte et al., 2008), an approach previously used to identify brain regions whose selectivity and coding properties are concordant or discordant depending on whether their response to a given set of stimuli is similar or dissimilar. Here, we apply a related similarity analysis not to the activity patterns themselves, but to the models that generate them and their co-fluctuations (see *Discussion*). The results of this cross-training prediction analysis are shown in Figure 4B, which gives the average prediction performance (correlation between actual and predicted values) for the different ordered pairs of trial categories: the classifier is trained on the first category and tested on the second one. As expected, the best cross-prediction performances were achieved when the training and the testing trial categories were the same (entries on the diagonal of the cross-training prediction matrix). However, despite a slight drop in performance, correlations between actual and predicted values remained significant for all other cross-prediction pairs (off-diagonal entries in Figure 4B).

Figure S3 presents how well the single-trial values of each individual element was predicted from the values of the others, for all trial categories confounded (top row), as well as for each category separately. As for all trial categories regrouped, the category-specific EC models achieved good prediction performances. The values of the cross-validated correlations between the actual and the predicted values and their patterns of variation over the different individual oscillatory elements were highly similar across trial categories.

In conclusion, all category-specific EC models could reliably predict single-trial fluctuations of the oscillatory elements for any other trial category, suggesting that they shared a backbone of common predictive links. Still, the best predictions were achieved when the training and the testing categories were the same, meaning that the category-specific EC networks and associate hyperplanes of co-fluctuation also presented distinguishing traits. In other words, the internal interdependence structure of the single-trial fluctuating oscillatory portraits was subtly and adeptly tuned in response to the changes in the task conditions.

### Oscillatory portraits reliably discriminate task conditions

The EC networks computed for the different trial categories (Figure 4A) achieved the best (cross-validated) prediction when the training and the testing categories were the same, indicating trial category specific distinct features despite their similarities (Figure 4B). This suggests that single-trial portraits may include the fingerprints of different statistical generative models, through which trial categories can be discriminated from single-trial observations. As represented schematically in Figure 5A, we constructed supervised linear mixed models to pairwise discriminate between trial categories. We compared the cross-validated generalization performance achieved by classifiers using oscillatory portraits as input (left), with classifiers using oscillatory elements (right; see *Materials and Methods*).

Linear classifiers output continuous-valued probabilities indicating the accuracy of the classification given the provided input information. To decide whether a trial belongs to either one of the categories, we compared the output probability with an arbitrary decision threshold. This situation is typical in pairwise-discrimination paradigms and is usually addressed by evaluating performance in terms of a *Receiver Operating Characteristic* (ROC) curve analysis (Green & Swets, 1966). In this analysis, the decision threshold is systematically varied from a minimum to a maximum value, and the corresponding fractions of misclassified false positives (FP) and correctly classified true positives (TP) trials are plotted as parametric curves. Random-like decisions lead to ROC curves sitting along the diagonal of the FP and TP rates plane (as if the decision was taken by tossing an unbiased coin). However, any significant displacement of the curve toward the upper left corner of the FP and TP rates indicates better-than-chance discrimination, so that the Area Under the ROC Curve (AUC) can be taken as an overall quantification of performance across the spectrum of possible decision thresholds (see *Material and Methods*).

Figure 5B presents the AUC scores for trial-category separations achieved by classifiers based on oscillatory portraits (Figure 5A, left). Discrimination performance (AUC scores) varied across pairs of trial categories. Nevertheless, in all cases, linear classifiers operating on single-trial oscillatory portraits achieved above chance level discrimination (Figure 5B). We also assessed the trial-category discriminations performed by linear classifiers based on individual oscillatory elements vs classifiers based on oscillatory portraits (Figure 5A). The box plots in Figure 5B contrast the average AUC (over all pairwise separations) obtained based on oscillatory portraits versus individual oscillatory elements. The AUC for the portrait-based classifiers (median: 0.62) were significantly (two-sided sample t-test, 50.05, p<0.0001) higher than those for the individual element-based classifiers (median: 0.52). Figure 5C presents results for the discrimination between *NR-Miss* and *RS-Hit* trials, which was the best achieved pairwise trial-category separation. The performance by the portrait-based classifier is shown along with the one obtained with the model based on the most discriminative element < IC4, *Dur*, β >. The ROC curves for the two classifiers can be compared to the chance-level ROC curves obtained when trial-category labels are shuffled.

Figure 5 shows that trial-categories can be discriminated based on the information conveyed by oscillatory portraits but does not tell *how* the trained portrait-based classifiers manage to extract this information. One can get some insight by inspecting the coefficients of the trained classifiers. For a given pairwise discrimination case, different portrait elements will weigh differently; that is, their fluctuations will affect the classifier decision output differently. Furthermore, some of the coefficients are positive and other negative, indicating the direction of the influence on the fluctuations of each given element. The coefficients of all the fitted pairwise discrimination linear classifiers are summarized in Figure S4. These coefficients specify the orientation of a- hyperplane separating the typical subspaces of fluctuation for portraits of the two trial categories to separate. The sign and magnitude of these coefficients provide some information on the importance of individual elements in inferring the trial category, and on the direction of their relative variation between the two discriminated conditions. For instance, element <IC4, *Dur*, *β>* as also shown in Fig 5C, had a strong negative coefficient for the classifier discriminating category *NR-Miss* and *RS-Hit*, corresponding to the fact that oscillation <IC4, *Dur*, *β>* was smaller in trials of the *RS-Hit* type than of the *NR-Miss* type. Similarly, <IC1, *Dur*, θ> also showed a significant negative coefficient indicating that frontal theta power during movement was lower for *RS-Hit* as compared *NR-Miss* as supported by literature that frontal theta increases with sensorimotor prediction and kinematic error (Arrighi et al., 2016). Consecutively, <IC1, *Dur*, θ> showed a strong positive coefficient for all comparisons against *Ct-Miss* suggesting that frontal theta power was higher for *Ct-Miss* compared to all trial categories and this is consistent with the observation that the subjects experienced the highest error in *Ct-Miss*. However, the interpretation of these coefficients was not always as easy, as coefficients themselves are degenerate. The same discrimination indeed could be performed by classifiers with different coefficients if they specify the same separation hyperplane. As previously shown, oscillatory elements were inter-dependent (Figures 3 and 4), therefore their values could be expressed as linear combinations of other elements. By replacing within the discriminating linear model an element variable by a corresponding linear combination of other elements variables, one would obtain by construction an equivalent classifier, but with different coefficients. This degeneration of classifier coefficients serves as a reminder that the discrimination between trial categories is always performed in terms of the entire high-dimensional oscillatory portrait, even when some individual coefficients are larger than others in a specific instance of implemented model.

### Oscillatory portraits predict intertrial behavioral variations

We have demonstrated that single-trial oscillatory portraits (in contrast to single-trial elements) carry information sufficient to discriminate trial categories reliably. Individual trials differed also in fine details of the movement kinematics, such as precise movement error and duration, since, even within a specific trial category, such features could fluctuate from trial-to-trial. We asked therefore whether fluctuations of oscillatory portraits could also predict these detailed fluctuations of behavioral features, beyond a discrimination between categories of behavior.

For this aim, we used linear mixed models (see *Materials and Methods*) receiving as input single-trial oscillatory portraits and producing as output continuous-valued estimates of trial-by-trial movement error or movement duration (Figure 6A; see *Materials and Methods* for exact definitions of the two behavioral features). Prediction performance was quantified by the Pearson’s correlation between the actual and predicted values. Figure 6B presents the performances achieved for predicting movement error and duration for all trial categories confounded, whereas Figure 6C (top) provides the detail of the correlation values for the different trial categories separately. Figure 6C (bottom) shows representative scatter plots of predicted vs actual movement error and duration values for the trial category *NR-Hit*. For all categories of trials, the correlations between the actual and the predicted values were strongly significant and the mean squared error was significantly less as compared to their shuffled versions for both movement error and movement duration of all trial categories (all are listed in Table 1). The correlations were also higher than those obtained by using linear models based on individual oscillatory elements (Figure 6B). The best prediction for movement error was achieved for the *Ct-Miss* (0.51) and *RS-Hit* (0.47) trials, whereas the best prediction for movement duration for the *Ct-Miss* (0.39) and *RS-Hit* (0.39) trials.

**Figure 6:**
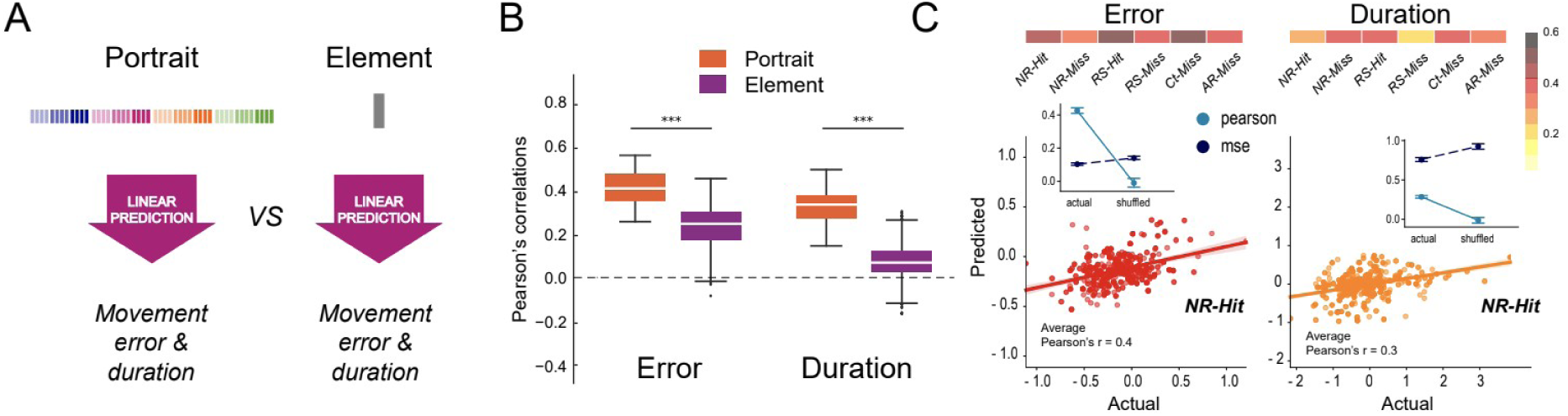
Inter-trial fluctuations in movement kinematics are reliably predicted from oscillatory portraits. **A)** Classifiers based on oscillatory portraits vs individual elements were trained to predict single-trial values of movement error or duration. **B)** Prediction performances are measured as the Pearson’s correlations between the actual and predicted values. The results are shown for all trial categories pooled together. The portrait-based classifiers outperformed the element-based classifiers for both movement error and duration. **C)** Pearson’s correlations between actual and predicted movement error and duration for each trial category. For the example trial category (NR-Hit), scatter plots of actual vs predicted values of movement duration or error. The colors of the scatter plots correspond to the correlation colormap in (C). The insets show the Pearson correlation and mean squared error over multiple folds between actual and shuffled versions of the data.

**Table 1:** Pearson’s correlation between actual and predicted values for movement error and duration in Figure 6C. All correlations are significant (p <0.0001), given values are averages over 20 folds with 75% as training and 25% as testing data.

| Trial category | Behavioral feature |  |
| --- | --- | --- |
|  | Movement error | Movement duration |
|  | Pearson's correlation | Pearson's correlation |
| NR-Hit | 0.42 | 0.28 |
| NR-Miss | 0.3 | 0.37 |
| RS-Hit | 0.47 | 0.39 |
| RS-Miss | 0.38 | 0.2 |
| Ct-Miss | 0.51 | 0.39 |
| AR-Miss | 0.35 | 0.35 |

The fact that portrait-based regression models achieved high and significant correlations for all trial categories taken separately demonstrates that the inter-trial behavioral fluctuations within each trial category could be captured, and not only the broad average kinematics differences produced by the manipulation of the task condition. A possible explanation for this successful prediction is that a tight coupling exists between fluctuations of behavior, on one side, and fluctuations of oscillatory portraits over trial category-specific manifolds, on the other. To probe the plausibility of this hypothesis, EC networks must be extended to encompass influences from and to motor behavior itself.

### Widely distributed and dichotomous coupling between movement error/duration and oscillatory portraits

The observations we have just described suggest possible mutual directed influences between the oscillatory portraits and the movement kinematic features. We generalized therefore our EC analysis (so far applied to oscillatory elements only) by including behavioral measures (movement error and duration) as additional nodes in the EC networks. The behavior-augmented EC networks for the different trial categories are presented in Figure 7A. As in Figure 4, only strong (see *Materials and Methods*) links are shown and colored according to their source node. Various observations can be made about the organization of these networks.

**Figure 7:**
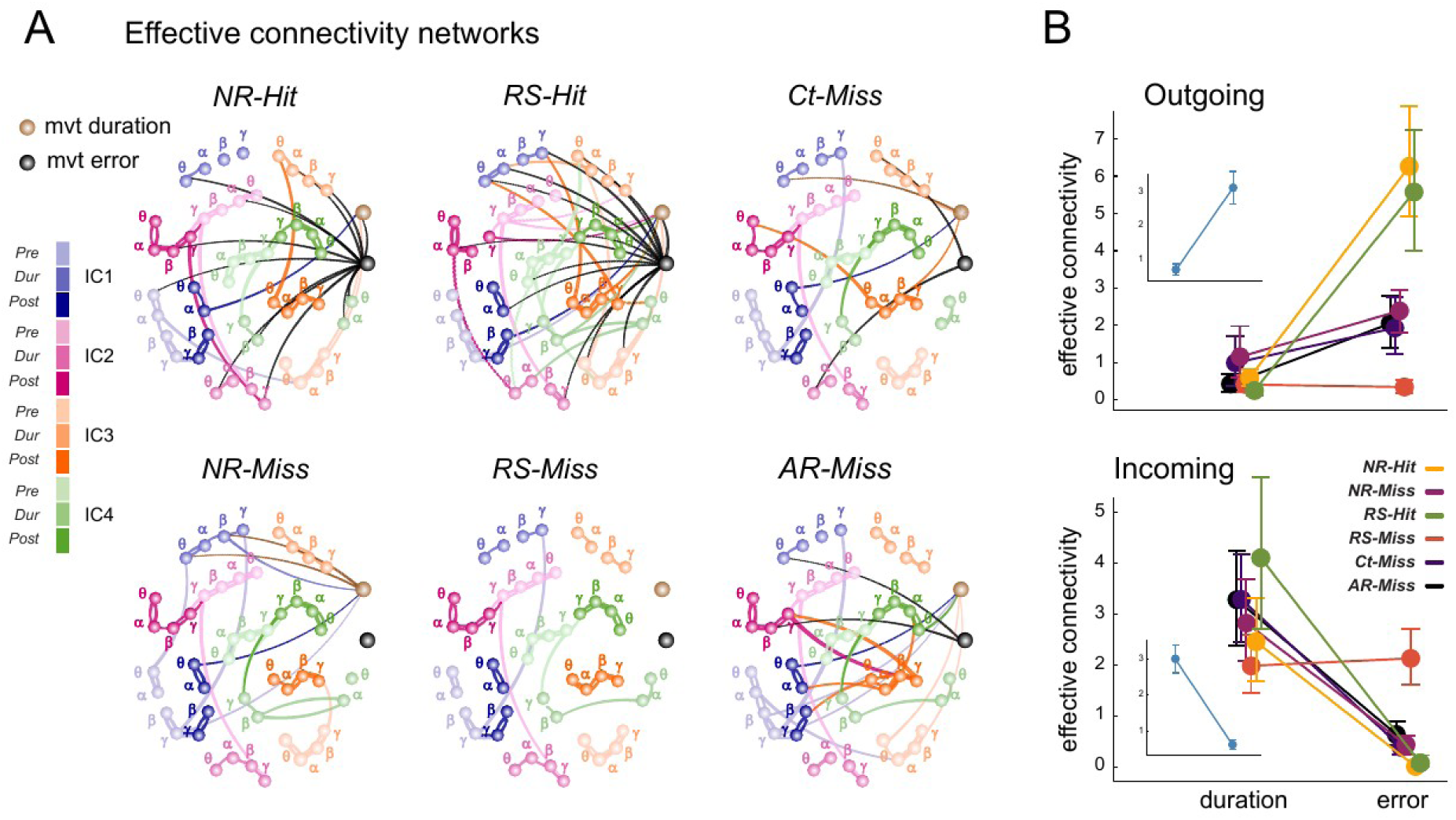
Effective connectivity networks ascribed movement error and duration as bottom-up and top-down features, respectively. **A)** Graph representation of effective connectivity integrating movement kinematics measures as additional nodes. The error and duration nodes are represented in black and brown, respectively. Only significant links are presented and colored according to their source node. **B)** Total in- and out-strengths of effective connectivity originating from behavior-related nodes (i.e. some of weights of EC connections) for all trial categories pooled together. The insets show the results for all trial categories pooled together. Caps mark the 95% CI of the distributions. Top, outgoing projections from the error node are significantly stronger than those from the duration node. Bottom, incoming projections from the oscillatory elements to the duration node are significantly higher than those to the error node.

First, the movement error node –and the movement duration node to the lesser extent– rather than being strongly coupled to only a few specialized hub nodes, sent diffuse coupling links to many oscillatory element nodes. For instance, as shown in Table S1, in the *RS-hit* condition, the movement error node sent one third of its connections to IC1 nodes, one third to IC3 nodes, and the remaining third to IC2 and IC4 nodes, displaying poor spatial preference. Analogously, in the *NR-hit* trials, movement error outgoing connections were spread across frequencies, with ∼36% of them reaching theta nodes, ∼27% alpha nodes and the remaining fraction equally split between beta and gamma nodes. Interestingly, both *Hit* trials showed poor spatial, spectral and temporal preferences, i.e. showed widespread connections across brain areas, frequencies and movement times.

Second, behavior-related nodes exhibit strong asymmetries between their outgoing and incoming connectivity. The movement error node influences the oscillatory portraits more than the oscillatory portrait influences the movement error, as revealed by a total out-strength (i.e. sum of all outgoing couplings; Figure 7B top) much larger than its total in-degree (i.e. sum of all incoming couplings; Figure 7B bottom, number of outgoing vs incoming connections are shown in Figure S5). The movement error node emanates the largest numbers of projections for *NR-Hit* and *RS-Hit* trial categories (Figure S5). The situation is inverted for the movement duration node, which is influenced by the oscillatory portraits more than vice versa, as revealed by an in-strength larger than the out-strength. The movement duration node receives most projections for *RS-Hit* and *AR-Miss* trial categories (Figure S5, T2).

Third, the connectivity of behavior-related nodes with oscillatory portraits is characteristically modulated by the trial category. Figure 7B shows in-strengths and out-strengths separated by trial categories (see the insets for all-trials pooled). During *Hit* trials (both *NR-Hit* and *RS-Hit* trials) the movement error maximally influences oscillatory portrait fluctuations (boosted out-degree) while the influence is substantially reduced during *Miss* trials, reaching a minimum for *RS-Miss* trials. Therefore, larger integration of information about movement error by the oscillatory element fluctuations is associated with successful behavior, which could possibly reveal, on the other side, a failure to integrate movement error information associated with unsuccessful behavior; *Miss* trials (see *Discussion*).

We also quantified the distribution of outgoing projections from the movement error node according to the spatial, temporal and spectral components (Table S1 and T2). Interestingly in *Ct-Miss* trial category which represents a large, unexpected kinematic error, movement error “regulates” predominantly the parietal and sensorimotor spatial component, notably, the θ spectral component in *Ct-Miss* in agreement with Arrighi et al. (2016). Another noteworthy observation was that the two *Hit* trial categories, *RS-Hit* and *NR-Hit,* not only had the highest number of outgoing projections (Tables S1 and S2), but also were more heterogeneous in their spatial and spectral targets, which supports our hypothesis that a successful behavior reflects greater integration of error information in the underlying network.

## Discussion

Inter-trial power variability is not “just noise” but carries relevant information. This had already been shown by studies that successfully related behavioral and electrophysiological fluctuations at single-trial level (Cohen & Cavanagh 2011; Torrecillos et al., 2018; Lofredi et al., 2019). However, most of these studies focused on individual oscillatory processes at a time, with the aim of mapping what we called here “oscillatory elements” to specific computations. Here, we move from a different tenet, hypothesizing that oscillatory element in different brain regions, trial epochs and frequency bands do not occur independently from each other, but instead are closely and dynamically coordinated, and that studying them separately provides only fractionated views of the overall system’s operation in relation to the task performed.

By considering high-dimensional oscillatory portraits, instead of basic oscillatory elements, we significantly improved: (1) trial-category pairwise separation; and (2) prediction of single-trial movement kinematics (movement error and duration). We propose that this advancement is not only methodological; but also theoretical, as it is consistent with the idea that behavior is not controlled by a myriad of unrelated oscillatory processes, but instead by collective dynamical modes that manifest themselves exerting distributed co-modulations of oscillations at different locations (Atasoy et al., 2016; Kirst et al., 2016). We modeled here these inter-dependencies as networks of Effective Connectivity (EC) that were neither totally disconnected –revealing fully segregated oscillatory elements–, nor completely connected –revealing global integration–, but had a homophilic organization between order and disorder, providing a mixture of integration and segregation which has been denoted as “complexity” (Tononi et al., 1998) and which is necessary to support rich system level emergent computation (Crutchfield, 2011). Such effective connectivity may in part stem from intrinsic constraints because of collective dynamics partially shaped by the underlying structural connectome (Honey et al., 2007), but not completely shaped by it because of nonlinearities and self-organization (Battaglia et al., 2012).

The importance of anatomical structure in shaping the EC was confirmed by marked degree of spatial homophily, indicative of long-range correlations coexisting with stronger local interactions. Frequency and time homophily are above chance level as well. On one side, frequency homophily in EC reflects coordinated fluctuations at the same frequency across different regions and across all epochs and it may thus denote the coexistence of multiplexed communication-through-coherence channels (Vezoli et al., 2021). However, our EC measures statistical relations between power fluctuations rather than actual phase-coherence and hence we cannot confirm this hypothesis. On the other side, temporal homophily manifests a large degree of simultaneity in the fluctuations of all oscillatory elements in the same time-range, irrespective of their frequency and location. This may indicate that high broadband power boosting events tended to simultaneously occur across all regions, possibly linked to slow modulations by non-neural physiological processes (Yuan et al., 2013) or global activity patterns linked to fluctuating arousal (Raut et al., 2021). Another possible explanation of coordinated spectral changes across all frequencies is that the spectrum was globally modified, in all its aperiodic and periodic components, by physiological processes as alterations of the E-I (excitation-inhibition) balance (Gao et. al, 2017). Space, frequency and time homophilic connections did not represent however 100% of total connections, i.e. some non-homophilic connections still existed. Nonlinear dynamics indeed can coordinate activity of very distant regions, especially when correlation lengths and susceptibility diverge in proximity of a critical point (Byrne et al., 2022) thus reducing space specificity. Furthermore, it can smooth frequency specificity by mediating cross-frequency influences (Breakspear & Terry, 2002; Kasatkin et al., 2017; Dellavale et al., 2020). Finally, baseline fluctuations can interact with presented stimuli in a non-additive way (He, 2013; Wainio-Theberge et al., 2021) so to affect the fluctuations of activity in temporally consecutive epochs and blur time-specificity of EC.

We did not explore here intrinsic connectivity in the resting state, but other studies have shown that, even at rest, oscillatory elements are widely coordinated (Mostame et al., 2021). Nevertheless, we revealed substantial similarity between EC networks across different trial categories, hinting at a possible intrinsic origin of the skeleton of homophilic connections that all these EC network share. We also propose that the detailed topology of EC was flexibly adapted around this shared scaffold to fine-tune to different behavioral demands (Figure 4) and as an effect of the integration of movement error information (Figure 7). Another possibility is that EC is preserved identically across all trial categories and that the trial-category specificity visible in Figure 4 was just a manifestation of overfitting. This is unlikely, however, as we determined our EC models using a cross-validation procedure that was robust against overfitting. Therefore, the observed EC and its modulations by the task constraints may reflect the fact that neural activity fluctuations are constrained to sample lower-dimensional manifolds within the higher-dimensional space of possible dynamic configurations (Gao & Ganguli, 2015; Gallego et al., 2017; Chauduri et al., 2019; Queralt et al., 2021), as an effect of dynamic and structural constraints, and that these manifolds are slightly deformed by behavior-related steering inputs. The manifold notion provides a geometric interpretation for the emergence of interdependence constraints. A point bound to move on a circle in a 2D plane, cannot perform independent excursions along the horizontal or the vertical axes, as the need to remain tangent to the circular contour entails correlations in the vertical and horizontal displacements. Analogously, the fluctuations of two oscillatory elements (e.g. the betas in two different regions) will be correlated by the need to sample a manifold prescribed by the current dynamical mode. The manifolds constraining the co-fluctuations of oscillatory elements may be slightly morphed by sensory or endogenous biasing inputs steering the system’s configuration as a whole, rather than individual elements so to distort the manifolds over which system’s trajectories unroll. The directions locally tangent to the dynamic manifold would be thus different for different trial categories, and a classifier could learn to discriminate them, as we showed in Figure 5.

Beyond clear behavioral differences between trial categories, we also found that fine fluctuations of portraits on top of their trial category specific manifold were inter-related with fine trial-to-trial fluctuations in behavior itself. Indeed, even within a same category of trials, different trials may differ in their exact values of movement error and movement duration, and we showed that these fine variations as well could be predicted significantly from portraits (Figure 6). By augmenting EC networks to encompass as well directed influences from and to these behavioral features, we could identify very distinct dominant directions of coupling for movement error and duration. The duration-node was mostly subjected as a target of influences from the oscillatory elements, whereas the error-node acted mostly as a source of influence on the oscillatory elements. The dominant direction of the EC between movement error and the oscillatory elements fits with the idea that coordinated oscillations implement a movement monitoring system. Interestingly, the EC from the error movement node toward the oscillatory portraits was the strongest in the trials in which participants moved accurately enough to successfully shoot the target (*NR-Hit* or *RS-Hit* trials). In these trial categories, the projections from movement error to the oscillatory portraits were also more heterogenous across spatial and spectral components. This finding may reflect that, in these categories of trials, oscillatory fluctuations were more precisely modulated by error leading thus to more accurate movement. In contrast, this influence was considerably weaker in trials that were performed in the same conditions (no or an expected visual rotation), but in which participants missed the target (*NR-Miss* and *RS-Miss*). This may suggest that failure to integrate movement error information may cause –rather than just correlate with– movement trajectories missing the target.

The behavioral feature that is monitored depends on the experimental paradigm. Our task was designed so that slight deviations in movement direction were enough to miss the target, and the subject were instructed to focus on meeting the target. This may explain why lower prediction accuracy was observed for movement duration than for movement-direction error. Also, with our task design, movement duration was extremely controlled and was relatively less fluctuating than movement error. It would be interesting to conduct the same network analyses on data collected using an experimental paradigm where movement duration, and not movement error, is the critical kinematic parameter for task success. We may, in this case, observe similar monitoring connections for movement duration.

The influence of the movement-error node could spread over a multiplicity of different elements; the oscillatory coordination manifold itself –and thus the EC network as a whole– was modulated by the injection of information about movement error. The fact that the integration of behavior-related information shapes a manifold of oscillatory coordination rather than steering individual oscillatory elements may have an important advantage: the system’s dynamics remain, at least in part, unconstrained, if specific co-fluctuation patterns are respected. As such, the large residual variability of system’s instantaneous configuration may serve as the device through which statistical inferences underlying sensorimotor control are neurally implemented, under the joint influence of prior expectations –encoded in baseline fluctuating activity– and external evidence (Körding & Wolpert, 2006). Future studies could test this hypothesis by observing how the EC between portraits and behavioral features evolve to get progressively more structured through learning.

Finally, our “effective connectivity” approach has several limitations. A variety of effective connectivity notions have already been proposed in the literature, with various alternative definitions, ranging from generic directed forms of functional connectivity (Sporns, 2007; Battaglia et al., 2012) to descriptions of causal inter-relations based on models of varying complexity and degrees of abstraction (Aertsen et al., 1989; Friston et al., 1994; Gilson et al., 2020). The EC notion we propose here is based on a simple and completely linear mixed model formulation. Unlike more usual EC frameworks which do fit models directly on time-series of neural activity, we chose here to operate directly on tables of band-integrated power values. Our aim indeed was not so much to identify causal influences between distinct neural populations and brain regions, as in classic formulations of EC, but rather to highlight statistical interdependencies between neural features usually studied as if they were independent.

A second limitation is linked to the fact that we define oscillatory components averaging power over broad generic bands rather than tailoring the bands to subject-specific peaks. However, using broad bands guarantees that these subject-specific peaks are included in the integration range, irrespectively from their exact location. Furthermore, we allow ourselves to capture as well information from aperiodic components of the spectrum, which, although far from the main spectral peaks, still convey a great lore of physiologically relevant information useful to determine system’s state (Donoghue et al., 2020). Finally, instantaneous frequencies can fluctuate stochastically across time and trials (Xing et al., 2012, Feingold et al., 2015), so that focusing on excessively narrow frequency bands may lead to excluding out-of-peak oscillatory events from the analysis which may be weaker but still carry relevant information about behavior (Douchamps et al., 2022). It is possible that superior predictive performance could be achieved by a better characterization of fluctuating oscillatory bursts beyond simply broadly integrating over them.

A third limitation of our approach could be then its linearity, approximating manifolds of covariance as simple hyperplanes, while they could be generally non-linear and curved. In the future, topological data analyses approaches could be used to extract the more general topological structure of the observed point-clouds (Carlsson et al., 2005) in oscillatory portraits space and attempt superior prediction and discrimination based on robust and invariant metrics of topological differences.

To conclude, why do predictions based on portraits outperform predictions based on individual elements? We propose here that this superior performance stems from the fact that oscillatory elements are parts of a distributed oscillating neural system which is internally coordinated and collectively monitors and controls behavior. Future extensions of this work could explore this hypothesis further to check whether the superior performance is due to the combination of *unique* information that oscillatory elements separately convey; or to *redundancy* between elements, allowing to better separate signal from noise; or, yet, to their *synergy*, i.e. capacity to convey jointly information beyond the sum of the parts (Wibral et al., 2017). Detecting such synergies may provide indeed even stronger arguments in favor of a genuinely collective functioning of sensorimotor control systems.

## Materials and methods

### Participants

A total of 24 healthy adults (8 females) aged 26.5 years (range 20-32 years) took part in the study. All participants were right-handed, as assessed by the Edinburgh Handedness Inventory (Oldfield RC, 1971) and all had normal or corrected-to-normal vision. All participants were free of known neurological or psychiatric disorders and gave informed consent according to a protocol approved by the Ethics Board of the Aix-Marseille University. They received monetary compensation for their participation.

### Experimental setup

The experiment was performed using a robotic exoskeleton (KINARM, BKIN Technologies) that allows recording flexion and extension movements of the elbow and shoulder joints in the horizontal plane. The rotation of the visual feedback of the hand was applied using a semi-silvered mirror preventing direct vision of the hand. A cursor representing participants’ index fingertip and the visual display were projected onto the same plane as the (invisible) hand.

### Task

The task and the experimental protocol has been already described in detail by Jahani et al. (2020), who presented other results on the data recorded during the same experiment. Participants were required to make ballistic movements with no on-line corrections. The starting position was indicated by a 0.75cm diameter white circle located at the center of a large concentric blue ring (10 and 14cm radius for the inner and outer contour, respectively). Throughout the experiment, three possible targets located 5cm away from the starting position were indicated as 0.3cm diameter dark gray circles: 50°, 80° or 110° from the 0° straight-ahead direction (Figure 1A). To initiate a trial, participants had to maintain their index finger in the start circle for 2000ms, after which they were warned to get ready (*Ready* signal): the start circle disappeared, and the target was indicated (one of the three targets turned from a gray to a white circle). Following a 1500ms delay, the target was filled in white (“turned on”) indicating that the movement could be initiated (*Go* signal). Importantly, participants were clearly informed that they were not performing a reaction-time task and that they should take all the time they needed to prepare their movement. They were instructed to move through (“shoot”) the target without stopping and to end their movement between the inner and the outer contour of the concentric ring. They were also required to move fast enough so that their hand moved 5cm away from the start position within 250ms, computed from the time when its speed exceeded 5cm/s. Participants received visual feedback about their performance at the time the finger-tip cursor reached 5cm away from the starting position, hitting or missing the target: (1) the target exploded when the movement was fast and accurate enough (target hit); (2) the target turned red when the movement was fast enough but not accurate enough (target miss); (3) the target turned green when the movement was too slow, independent of its accuracy. According to their verbal reports, participants enjoyed the explosion of the target, which was experienced as rewarding. In order to avoid on-line movement corrections, the finger-tip cursor was turned off when the hand crossed the 10cm radius inner contour of the ring. Upon movement end, the arm was passively brought back by the robot to the start position. The finger-tip cursor and the starting-position circle reappeared only when the hand was back in its initial position. Each trial lasted about 7sec. Participants were asked to keep their eyes fixed on the aimed target throughout each trial.

### Experimental protocol

The experiment was made up of two sessions (familiarization and experimental) run on two different days, during which participants performed two categories of blocks: *Baseline* blocks comprising only unperturbed (no-rotation) trials and *Mixed* blocks in which a visual rotation (+30° or −30°) was applied in selected trials. In the Mixed blocks, short series of rotation trials alternated with no-rotation trials. The rotation-trial series counted 4 movements to the same target, whereas the number and targets of the no-rotation trials (*NR* trials) interleaved in between varied pseudo-randomly (at least 4 successive no-rotation trials) (Figure 1A). That is, participants could not predict when the rotation would be introduced. As a result, their hand trajectories were always clearly deviated and the target largely missed in the first trials of the rotation-trial series (*Ct* trials in Figure 1B). However, participants were informed (during the familiarization session) about the properties of the 4-rotation-trial series; that is, they knew the visual rotation would be applied in the three following trials as well (*RS* trials). Participants also knew that the rotation would be removed after 4 trials; that is, in the trials immediately following a 4-rotation-trial series (*AR* trials), they would have to quit the strategy and aim again directly at the target that was illuminated.

Each *Mixed* block comprised 18 rotation-trial series, and 96 no-rotation trials pseudo-randomly distributed in between, for a total of 168 trials. The direction of the rotation, clockwise (30°CW) or counterclockwise (30°CCW), applied in the rotation-trial series was kept constant throughout each *Mixed* block, but reversed for each new *Mixed* block. Half of the participants started with a 30°CW *Mixed* block, the other half with a 30°CCW *Mixed* block.

During the familiarization session, participants received verbal instructions about the general task requirements. They performed at least 4 blocks of 20 trials with no visual rotation, followed by a block in which, after 4 no rotation trials, the visual rotation (clockwise or counterclockwise, counterbalanced across participants) was unexpectedly introduced for 5 trials. After participants had experienced the visual rotation, the experimenter explained in detail the nature of the perturbation and how they could counter it by a strategy consisting in aiming at the (clockwise or counterclockwise) neighboring target (see Jahani et al., 2020). They performed two *Mixed* blocks, each followed by a 32-trial *Baseline* block (400 trials in total). EEG signals were not recorded during this session. During the experimental session, after a 64-trial *Baseline* block, participants performed four *Mixed* blocks, each followed by a 32-trial *Baseline* block (864 trials in total). EEG signals were recorded throughout the session. Between each block of trials (*Mixed* and *Baseline*) and after the 84^th^ trial of each *Mixed* block, a ∼2min break was allocated. The preliminary session lasted about 1h30min in total (including robot calibration) and the experimentation session (including EEG-electrode placement and location recording) lasted about 3h in total.

### Behavioral data recording

Angular position and velocity data of the motor resolvers were collected at 1000Hz. Signals were down-sampled offline to 100Hz, and then filtered with a 2nd order zero-phase-shift low-pass Butterworth filter (cut-off frequency of 10Hz). Hand position and velocity were calculated from these angular data. Kinematic data were analyzed using custom routines written in MATLAB (MathWorks). Trials in which the hand was not maintained stable enough in the start-position during the delay between the Ready and Go signals (tangential velocity > 6cm/s), or in which the movement was initiated before the Go signal, were excluded from the analyses (∼1% of trials). Movement onset was defined as the time when the tangential velocity exceeded 5cm/s. The movement offset corresponded to the time when tangential velocity fell below 5cm/s and remained below this value for at least 1500ms. To quantify kinematic errors, we computed the perpendicular deviation, from the straight line that connects the starting position to the target, at maximum velocity. This measure quantifies error in initial movement-direction (feedforward component). To collapse data from different Mixed blocks, with opposite visual rotations (30°CW vs 30°CCW), we set the signs of the PD-vel values so that hand-path deviations in the direction of the visual rotation corresponded to positive values. Previously, we conducted preliminary analyses to test for differences between the movement errors induced by the two rotation directions. Movement duration was also calculated. Trials that were performed too slowly (∼4%) were excluded from the analyses.

Note that, in our task, success was conditioned by accuracy in movement direction, and not in movement duration. Hence, in the current context, variations in movement direction constitute movement errors, whereas fluctuations in movement durations are not detrimental. Hence, we use “movement error” interchangeably with “movement-direction error”, unless specified.

### EEG data recording and preprocessing

EEG activity was recorded continuously at 1024Hz using a 64-channel Biosemi ActiveTwo system (BioSemi) referenced to the Common Mode Sense / Drive Right Leg (CMS/DRL) contact. Electrodes were embedded into an elastic cap and distributed over the scalp according to the extended 10-20 EEG system. The electrode offsets, the voltage differences between the CMS and each active electrode, were monitored to remain within ±20 μV. For each participant, electrode locations and nasion and preauricular points were recorded by an infrared camera (Rogue Research). Electro-oculographic (EOG) activity was recorded with surface electrodes placed near both outer canthi (saccades) as well as under and above the right orbit (blinks). EEG continuous signals were re-referenced to the average of all electrodes, filtered between 2-70Hz (Butterworth order 2) and down-sampled to 256Hz. Non-stereotypical artifacts (that cannot be captured by ICA; cf. Makeig et al., 1997; Delorme et al., 2007) were identified and rejected upon visual data screening. Further analyzes were run using the free and open-source software Fieldtrip (Oostenveld et al. 2011).

### Independent Components (ICs) identification

The preprocessed EEG signals were cut into time-segments extending from −3.5 to +3.5ms with respect to outcome feedback, which covered approximately the complete trials, slightly variable in duration. The epoched EEG data were then submitted to ICA (*runica* algorithm). Time-frequency analyses were performed on the time-courses of the independent components (ICs). Single-trial signals were transformed in the time-frequency domain by convolution with the complex Morlet’s wavelets characterized by the ratio f0/σf = 7, with f0 ranging from 2 to 50Hz by steps of 0.5Hz. In order to calculate the event-related changes in beta power, the raw power data was log-transformed and then normalized relative to the average power calculated over all trials, as no clear baseline period could be defined during our task (Tan et al. 2014, Torrecillos et al. 2015). For each participant and each time point (50ms bin), power was averaged over trials within a specific beta frequency band (individually selected; see below) and smoothed using a Gaussian Kernel with 7-time points (350ms) full-width at half maximum. Our goal was to identify for each participant four different ICs that would capture oscillatory activities from four functionally distinct cortical regions. First (IC1), the medial frontal cortex whose oscillatory activity is known to be sensitive to error and reward (ref.). Second (IC2), the medial parietal cortex, involved namely in high cognitive processes (ref.). Last (IC3 and IC4), the left and right sensorimotor cortices.

For the ICs selection, we proceeded in two steps (Alayrangues et al. 2019). First, we pre-selected ICs based on their topographies. For this step, we defined spatial regions of interest (ROIs); ICs that exhibited the largest weighting within one of these ROIs were pre-selected. To capture activity of the medial frontal cortex, we considered an ROI including electrodes F1-Fz-F2-FC1-FCz-FC2-C1-Cz-C2, for activity in the parietal medial region, we used an ROI encompassing electrodes C1-Cz-C2-CP1-CPz-CP2-P1-Pz-P2, and for the left and right sensorimotor regions, we used two ROI including electrodes C3-C5-CP1-CP3-CP5-P1-P3-P5 and C3-C5-CP1-CP3-CP5-P1-P3-P5, respectively. (For one participant, we selected an IC with maximum weighting at electrode FC3.) Then, in a second step, we examined the time-frequency representation of the time-courses of the pre-selected ICs to retain for each participant one IC of each category. For this step, within the trial-period going from 0 to 2.5 sec relative to outcome feedback, we examined the time-frequency representations of the time-courses of the pre-selected ICs computed over all trials. For each individual and category of ICs, we selected the IC (most of the time, only one IC per participant was preselected) exhibiting the largest power variance between 17 and 40 Hz.

### Oscillatory portraits

Oscillatory portraits were defined as triplets of average power over frequency bands, cortical regions and time windows. That is, average power in each trial was defined as a vector of 48 values – each representing one of 4 × 4 × 3 triplets e.g. (f,r,t), where f = frequency band, r = cortical region and t = movement phase (Figure 2B).

The fixed frequency bands were defined over ranges commonly used in the literature a) theta (4-8Hz) b) alpha (9-12Hz) c) beta (13-35Hz) d) gamma (36-60Hz)

Cortical regions: The independent components of four EEG hotspots were considered for analysis: IC1, corresponding to medial frontal; IC2, corresponding to medial parietal; IC3, corresponding to contralateral (left) sensorimotor cortex; and IC4, corresponding to ipsilateral (right) sensorimotor cortex (see Alayrangues et al. 2019; Jahani et al.).

Movement phases:

The average power was measured at three phases of movement:

- Pre-movement (−1.5 seconds before the movement onset). This phase represents the planning of the movement.
- During-movement. In trials locked to the hand movement, this phase lasts from 0 to trial and subject dependent movement duration and represents the movement execution.
- Post-movement. In trials locked to the hand movement, this phase lasts from the end of the movement duration to 1.5 seconds after the movement offset. Post-movement phase is said to be relevant for indicating prediction errors for motor learning.

### Linear mixed models

The trial-by-trial analysis was done by converting average power in every trial to oscillatory portraits as described in section *Oscillatory portraits*. These oscillatory portraits formed the predictor variables of the linear mixed model. The random effects were modeled as intercept with subject as categories i.e, every subject was fitted with a different estimate of intercept (random intercept model).

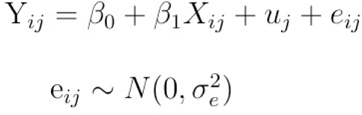

Where: Y_ij_ represented the predicted variable for subject j and trial number I; *β* _0_ the common intercept and *β* _1_ the common slope for all subjects; X _ij_ the matrix of k components for subject j and trial I; u_j_ the individual intercept for every subject; and e_ij_ the residual.

The package statsmodels was used to implement linear mixed models in python. The linear mixed model framework was used for the following analyses: deduce the effective connectivity network among the components and behavioral features; and pairwise separation of the trial categories.

### Pairwise separation of the trial categories

Within each trial category defined by the experimental protocol (NR, Ct, RS and AR trials - see Figure 1A) we distinguished between the trials in which participants successfully shot the target (Hit) from those in which they failed to do so (Miss). Participants missed the target in all Ct trials and in the vast majority of the AR trials. Thus, for our analyses, we considered the following 6 categories of trials: NR-Miss, NR-Hit, RS-Miss, RS-Hit, Ct-Miss, AR-Miss.

Linear mixed models were used to check if pairwise classification of the trial categories can be performed reliably using the oscillatory portraits. In order to check this, for all pairwise comparisons (15), the trial category label was the predicted variable i.e in the equation (1), Y _ij_ represented the trial category labels for a given trial category pair. The prediction was trained over 100 bootstrapped iterations of training and testing sets using a stratified shuffle split (sklearn). The measure used to gauge the accuracy of prediction was area under the curve (AUC) for region operating curves (ROC).. In order to check if the accuracy of the prediction was above the chance level, the same experiment was repeated with shuffled trial category labels.

To compare the efficacy of using portraits vs individual components in pairwise separation of the trial categories, separate classifiers were trained and tested either with portraits or an individual component. The accuracy distributions for all the task categories were pooled together and compared with a Mann-Whitney test.

### Cross training and testing paradigm

The cross training and testing paradigm consisted in: first, training the classifier for a trial category X; and, second, testing the performance of the classifier on all the other trial categories as well. The performance of the classifier trained and tested for the same trial category indicated how well a classifier captured the specificity of the trial category. The performance was measured as Pearson’s correlation coefficient between the actual and predicted values of the dependent variable (effective connectivity). The performance of the classifier trained for a trial category X but tested for the trial category Y indicated how well the classifier generalized over the trial categories X and Y, which was also an indirect measure of how similar the trial categories are.

The cross-training paradigm was used to quantify the generalization of the effective network across the task categories. The classifier was trained for a given task category and tested for the same as well as other task categories. The cross predictability was measured as the average Pearson’s correlation over the 48 components.

### Predict behavioral features

Generalized Linear Models (GLMs) were used to predict behavioral features such as movement duration and error, i.e. the predicted variable Y _ij_ represented the movement error/duration for the considered trial. The accuracy of prediction was measured as Pearson correlation between predicted and actual values as well as mean square error between the actual and predicted values. The prediction was done on 20 folds with 75% of data as training whereas 25% of data as testing data. The Pearson correlations and mean squared error were pooled for 20 folds and compared against the distributions for their shuffled counterparts (where the Y_ij_ was shuffled for training data).

### Correlation Matrix

Pairwise Pearson’s correlation coefficient between the oscillatory portraits (48 in number) was calculated for multiple bootstrapping iterations (50), consisting of randomly picked 75% of the data, and then averaged over these replicas. The mean correlation matrix is shown in Fig 3A.

### Effective connectivity

Linear mixed models were used to check if oscillatory portraits can be predicted from each other, i.e the predicted variable Y_ij_ represents one of the 48 oscillatory portraits and the rest 47 are the predictor variables. This analysis yielded an effective connectivity matrix among the oscillatory portraits and was done for all trial categories pooled together as well as separately. The effective connectivity matrices were calculated for multiple bootstrapping iterations (50), each time from 75% of the data randomly sampled. The effective connectivity was then averaged over the bootstrap replicas. The analysis was done for two versions. In a first one, the dependence analysis was restricted to oscillatory portraits only (without behavioral features). In a second one, the analysis was extended to include two behavioral features (movement error and duration) as a prediction targets as well as predictor variables. That is, we sought for the oscillatory portraits that significantly modulate the behavioral features and, as well, for the behavioral features that modulate the oscillatory portraits (with behavioral features). The accuracy of the prediction was measured by Pearson’s correlation between the predicted and the actual values.

### Graph representation

The effective connectivity network of oscillatory portraits from the section *Deduce effective connectivity* were converted into graphs for the ease of analysis and visual representation. Every oscillatory portrait/behavioral feature represented a node in the graph and dependence between the two portraits represents an edge. The graph representations were made using the python library networkx. The nodes and edges were arranged using a force directed spring algorithm (Fruchterman–Reingold), which arranges strongly connected nodes closer together than weakly connected nodes. However, for the ease of comparison, the arrangement of the spring algorithm was fixed for all trial categories, for a swifter visual comparison. The weights for all trial categories were pooled together to determine the threshold for “strong” vs “weak” connections. Only the weights larger than 97% percentile of the pooled weight distributions were shown as colored edges (strong) for visual clarity, whereas the rest are shown as gray edges (weak). We don’t represent edge weight via a different thickness of the plotted edge, i.e. all edges have the same thickness in the graphical representation. This explains why integrated strengths can sometimes be larger for graphs displaying visually fewer edge lines: there may be fewer significant edges, but they are stronger in weight (e.g. concerning the out-strengths of NR- and RS-hit EC graphs in Figure 7).

### Weighted Homophily

Homophily in a graph was defined as the probability to form a connection between two nodes sharing a similar feature, as compared to connections with other nodes having different values of the considered feature. The effective connectivity networks we investigated were weighted networks hence we estimated the average weight with which nodes connect with other nodes sharing a common label, of the temporal (movement phase), spectral (frequency) or spatial (brain areas) types, compared to the shuffled versions of the graphs, where the labels were randomly permuted across nodes. Such shuffling preserved the structure of the graph but disrupted the correlation of the temporal, spectral and spatial labels with the underlying connectivity.

The average spatial, spectral and temporal homophily was quantified as the relative percent ratio of the total weight of homophilic connections as compared to the total weight, i.e.

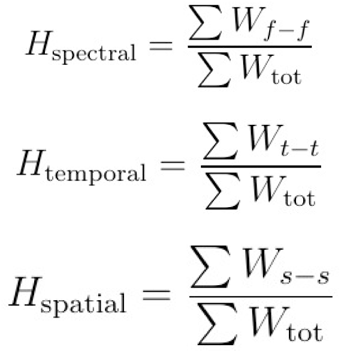

where W_f-f_ represents weights of spectrally homophilic links (such as theta-theta, alpha-alpha, beta-beta and gamma-gamma connections), W_t-t_ rweights of temporally homophilic node pairs (such as Pre-Pre, During-During and Post-Post), W_s-s_ represents weights of spatially homophilic node pairs (such as IC1-IC1, IC2-IC2, IC3-IC3 and IC4-IC4) and ∑ W_tot_ represents the total of all weights in the network.

Homophily was then defined, for each EC network, as a triplet consisting of the three homophily ratios:

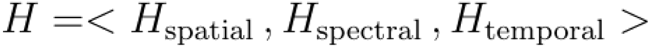

The % change in homophily ΔH for a trial category with respect to the others was calculated as a percentage change in the spatial, temporal and spectral homophily ratios for a trial category as compared to the values when all trials were pooled together.

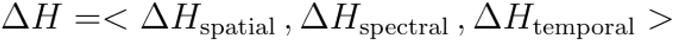

### Overlap between the distributions

Similarity between the distributions was calculated as a measure of distance in terms of Bhattacharya coefficient. The Bhattacharya coefficient between two probability distributions P and Q over the same data X is given as by:

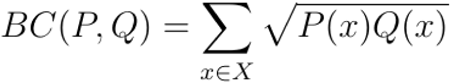

### Crossvalidation

Crossvalidation was performed by training the model for a section of data (training data) and then testing the model for the rest of the data (testing data), that the model did not encounter during the training session. The results for the testing data were then pooled over multiple iterations and an average performance was reported. Crossvalidation ensured that the results are not heavily influenced by outliers and represents consistent patterns over different sections of data. The crossvalidation was performed for all analyses, whenever possible, i.e.: when separating task categories (average performance); calculating effective connectivity; and, predicting behavioral features from the oscillatory portraits.

### Similarity of task categories to the pooled effective connectivity network

The similarity of the task categories to the pooled effective connectivity network (“all”) or to other trial-specific EC netwosrks was calculated as Pearson’s correlation coefficient between their weighed adjacency matrices flattened in vectors.

## Supplementary tables

**Table S1:** Break-up of outgoing connections from the movement error node to the oscillatory portraits.

| Trial category | Total number of outgoing connections (from Mvt. Error) | Spatial |  |  |  | Spectral |  |  |  | Temporal |  |  |
| --- | --- | --- | --- | --- | --- | --- | --- | --- | --- | --- | --- | --- |
| | | IC1 | IC2 | IC3 | IC4 | $\theta$ | $\alpha$ | $\beta$ | $\gamma$ | Pre | During | Post |
| AR-Miss | 2 | 50% | 50% | - | - | 50% | - | 50% | - | - | 50% | 50% |
| Ct-Miss | 2 | - | 50% | 50% | - | 100% | - | - | - | - | 100% | - |
| NR-Hit | 12 | 18.2 % | 36.4% | 18.2% | 27.3% | 36.4% | 27.3% | 18.2% | 18.2% | 18.2% | 72.7% | 9.1% |
| RS-Hit | 13 | 33.3 % | 16.7% | 33.3% | 16.7% | 33.3% | 25.0% | 8.3% | 33.3% | 25% | 50% | 25% |

**Table S2:** Break-up of incoming connections to the movement duration node from the oscillatory portraits.

| Trial category | Total number of incoming connections (to Mvt. Duration) | Spatial |  |  |  | Spectral |  |  |  | Temporal |  |  |
| --- | --- | --- | --- | --- | --- | --- | --- | --- | --- | --- | --- | --- |
| | | IC1 | IC2 | IC3 | IC4 | $\theta$ | $\alpha$ | $\beta$ | $\gamma$ | Pre | During | Post |
| AR-Miss | 6 | 50% | - | 33.3% | 16.7% | 50% | - | 33.3% | 16.7% | 66.7% | - | 33.3% |
| Ct-Miss | 3 | 33.3 % | - | 66.7% | - | 33.3% | - | 66.7% | - | - | 33.3% | 66.7% |
| NR-Hit | 4 | 33.3 % | - | 66.7% | - | - | 66.7% | 33.3% | - | 66.7% | - | 33.3% |
| NR-Miss | 3 | 100 % | - | - | - | 33.3% | - | 33.3% | 33.3% | 33.3% | 33.3% | 33.3% |
| RS-Hit | 10 | 22.2 % | 22.2% | 33.3% | 22.2% | - | 11.1% | 44.4% | 44.4% | 44.4% | 11.1% | 44.4% |

## Supplementary figures

**Figure S1:**
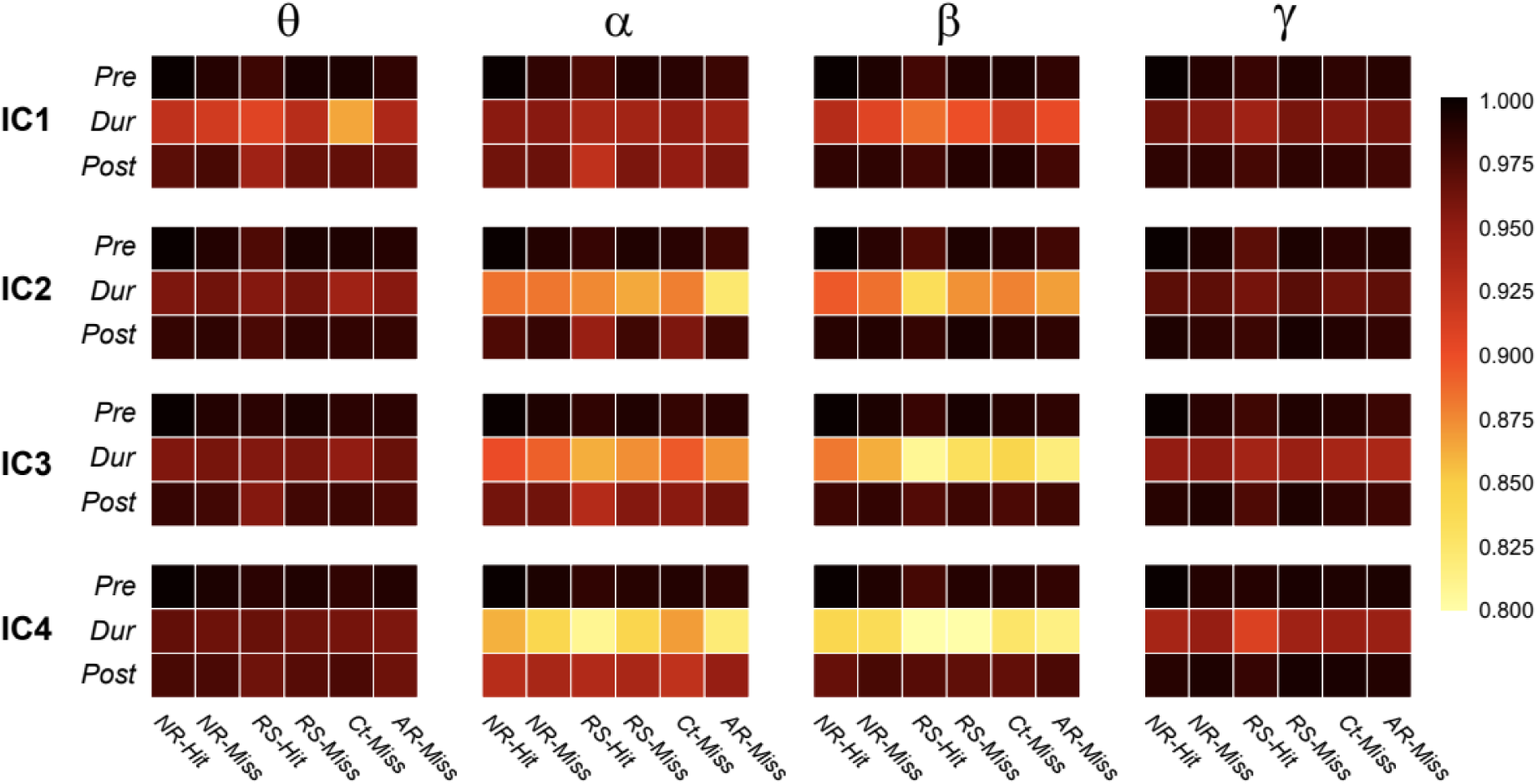
The distributions of the single-trial oscillatory element largely overlapped across the different trial categories. Element < IC1, Pre, θ > (top left cell) serves here as a reference and the density range overlap is measured as the Bhattacharya distance between the distributions.

**Figure S2:**
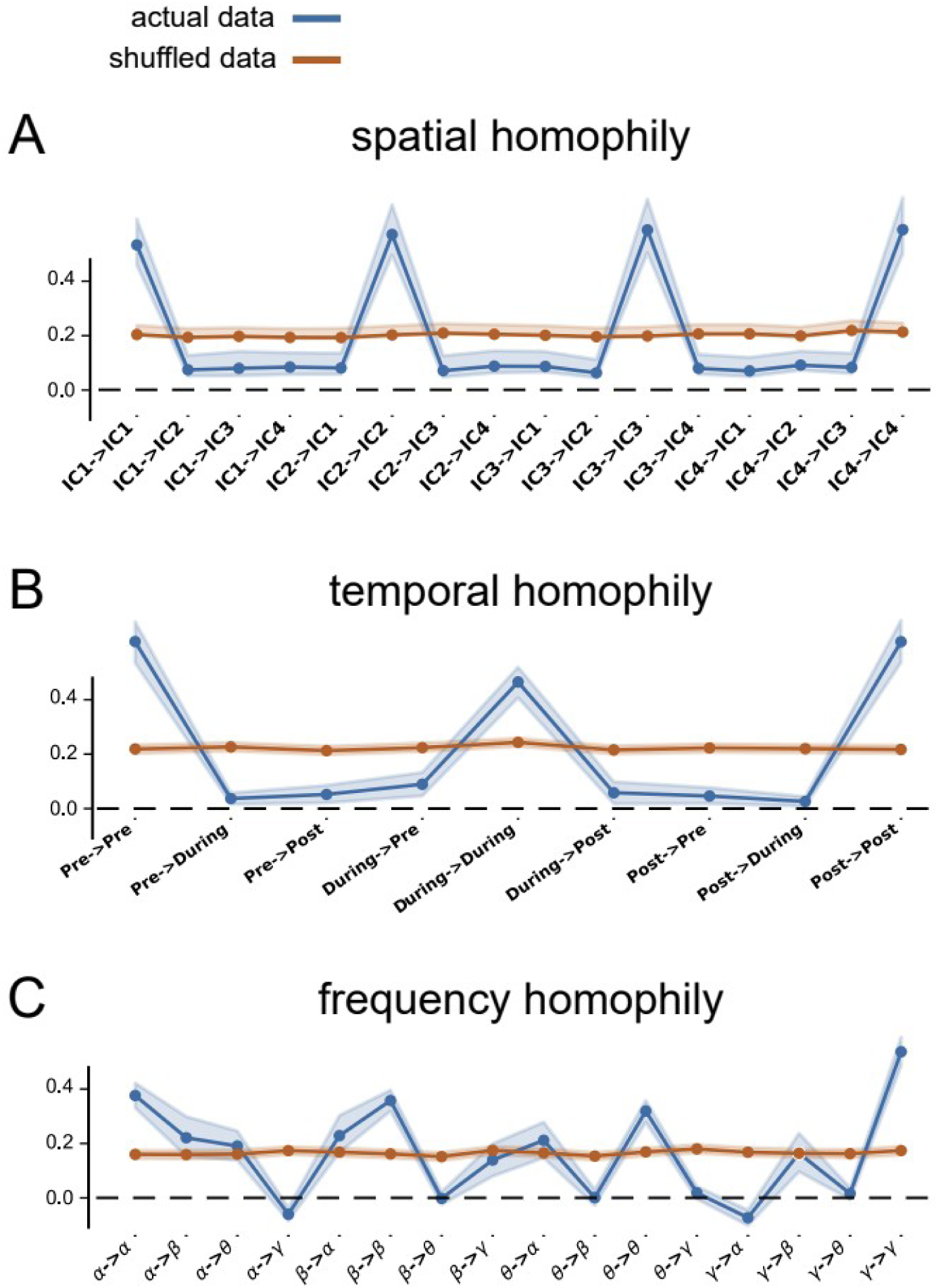
The effective connectivity (EC) network was highly homophilic. **A-C)** Average weight in the spatial, temporal and frequency homophily ratios, respectively. The combinations of node pairs are indicated on the x-axis. The average weights are plotted for the actual effective connectivity networks (blue) and their shuffled versions (orange). All the networks for all trials categories and subjects were pooled together. The spatial node pairs belonging to the same brain area (e.g IC1 <–> IC1) show increased average weight, which indicates that stronger effective connectivity between same spatial nodes irrespective of their temporal and spectral features. The shuffled version of the network shows no pattern with respect to spatial node pairs. We quantify the total degree of homophily as the ratio between the average of homophilic weights (i.e. the weights of links between nodes with same values of the considered label type) and the total weight. The degree of spatial homophily is thus evaluated to 73%. B and C) Same as A) but for temporal (85%) and spectral (62%) homophily rations, respectively.

**Figure S3:**
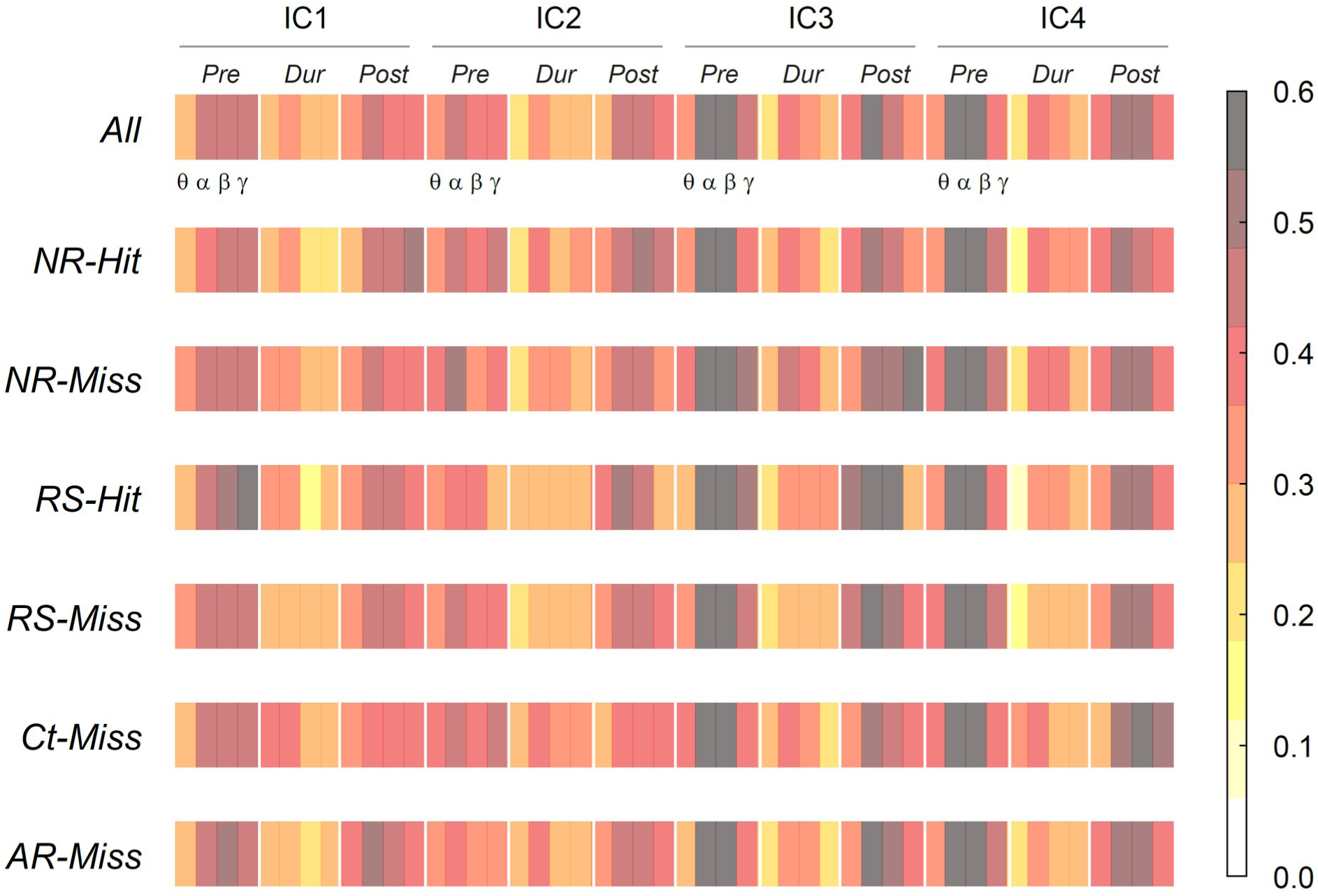
To generate effective connectivity (EC) networks, single-trial values of each individual oscillatory element were predicted from the values of the others. For each oscillatory element, the prediction performance was measured using Pearson’s correlation between the actual and the predicted values. This was performed for all the trial categories confounded (top row), as well as for the different trial categories separately.

**Figure S4:**
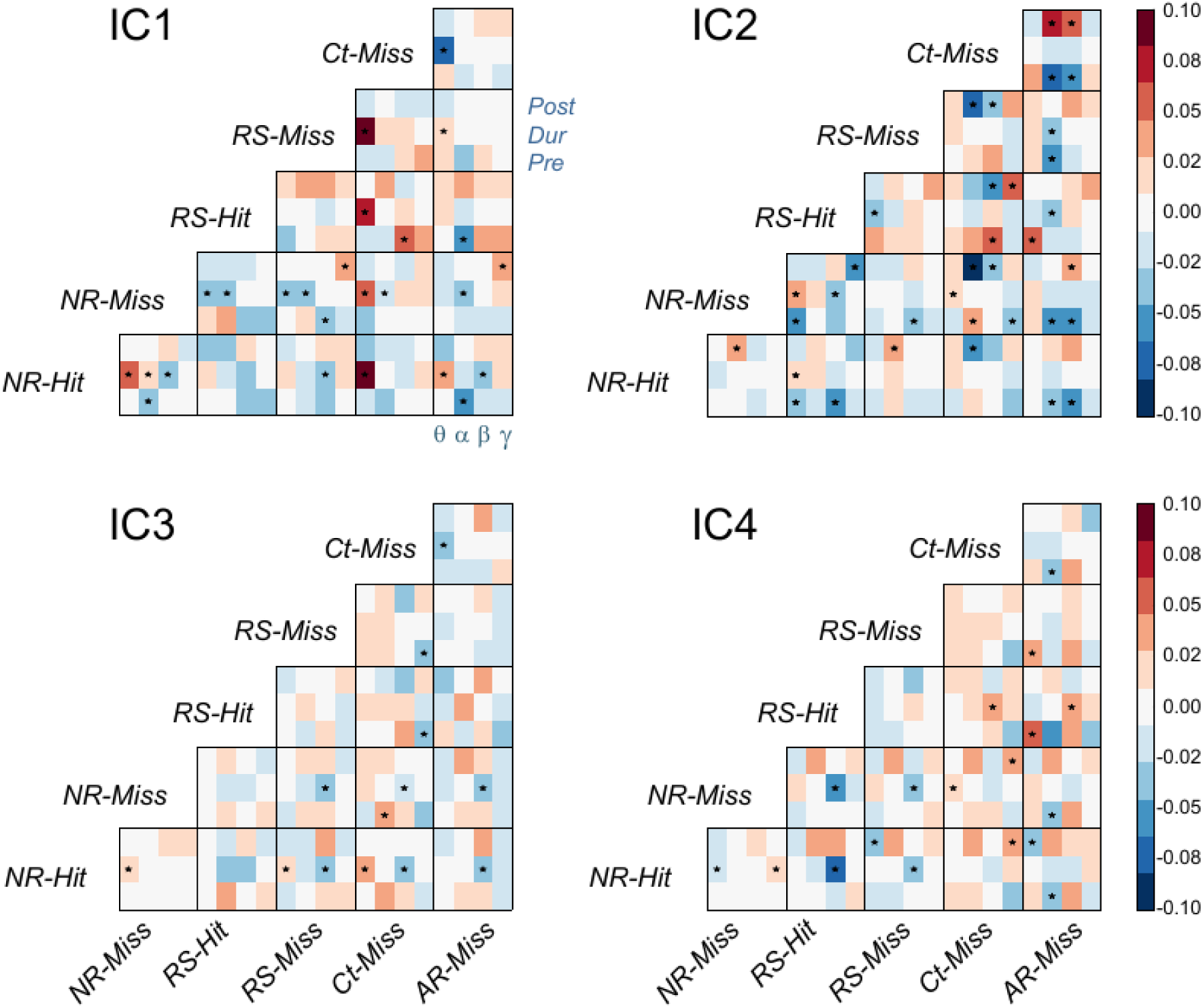
Coefficients of each oscillatory element for pairwise separation between the trial categories. The coefficients can take positive/negative values as indicated by the colors (warm = positive, cool = negative). Significant (p< 0.05) coefficients are marked with ‘*’. Negative(positive) values indicate that the considered element has smaller(larger) power for trials of the category in the row than for trials of the category in the column.

**Figure S5:**
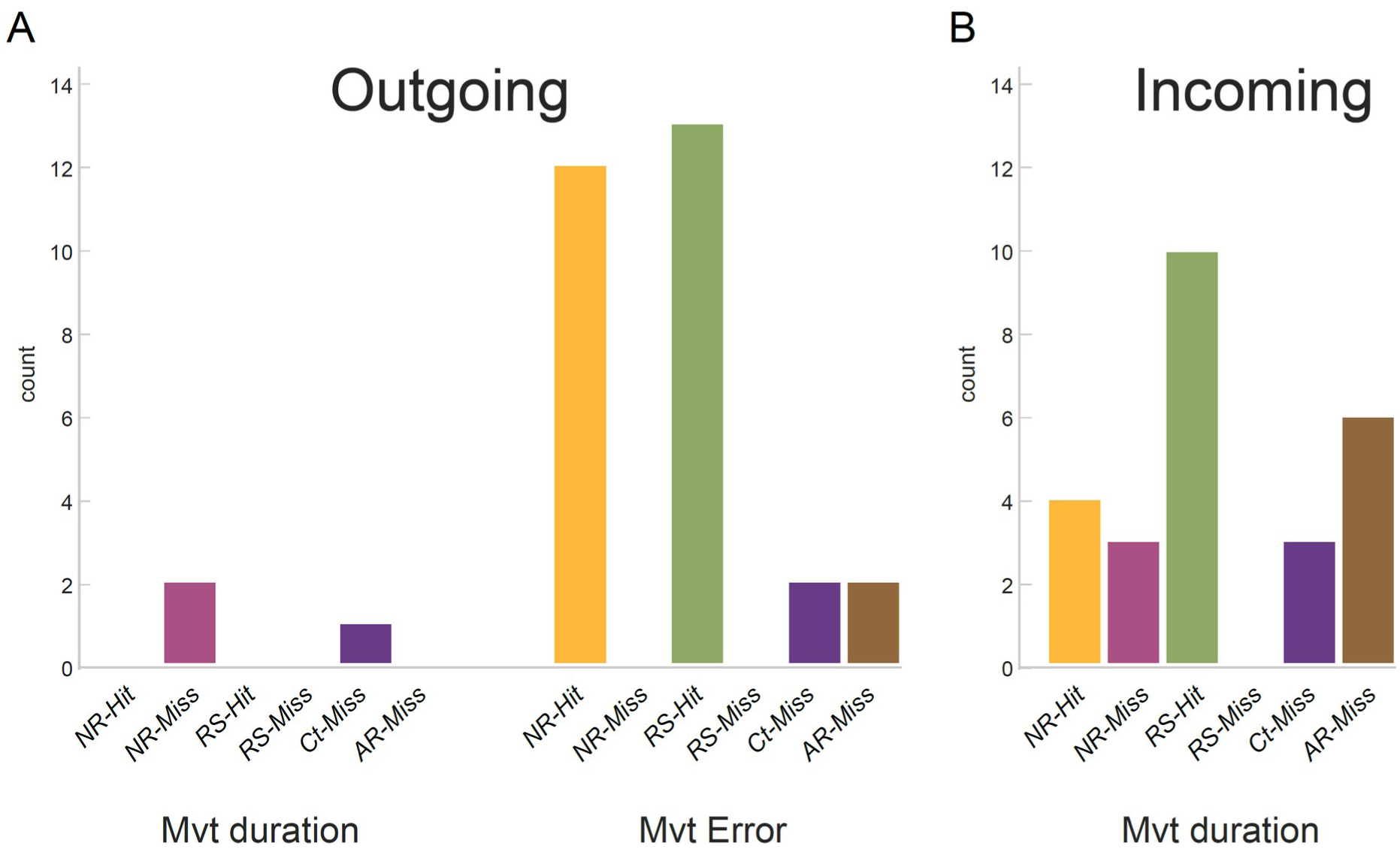
***A***) Number of outgoing connections from Movement Duration and Movement Error nodes in a thresholded network (only connections stronger than 94 percentile of weight distribution pooled over all trial categories were considered). Movement error, especially RS-Hit and NR-Hit have high number of outgoing connections. **B)** Movement error in any trial category had any strong incoming projections. Whereas Movement duration nodes have high number of incoming projections, especially for RS-Hit and AR-Miss. The breakdown of these connections are listed in Table S1 and S2.

